# Some new *Haliclona* species (Demospongiae, Haplosclerida) from British Columbia Shallow Waters and a Re-Description of *Haliclona mollis* (Lambe, 1893)

**DOI:** 10.1101/2024.04.03.587985

**Authors:** Bruce Ott, Neil McDaniel, Rick Harbo, Hugh MacIntosh

**Affiliations:** Bruce Ott, 4577 W 15 Ave, Vancouver, BC Canada V6R 3E8; McDaniel Marine Surveys, Vancouver, BC, Canada.; Research Associate, Royal British Columbia Museum, Victoria, British Columbia, Canada; Invertebrate Zoology, Royal British Columbia Museum, Victoria, British Columbia, Canada

**Author notes:** Correspondence author: Bruce Ott, 4577 W 15 Ave, Vancouver, BC Canada V6R 3E8, https://orcid.org/0009-0006-1624-5995. NOTE: THIS MANUSCRIPT IS NOT A PUBLICATION WITH RESPECT TO NEW SPECIES BUT A PRE-PRINT UNREVIEWED DRAFT.

**Keywords:** Porifera, Demospongiae, Haplosclerida, *Haliclona* (*Flagellia*), *Haliclona* (*Gellius*), *Haliclona* (*Haliclona*), *Haliclona* (*Reniera*), *Haliclona* (*Rhizoniera*), invertebrate systematics, British Columbia, marine biogeography

## Abstract

**Background:** A numer of *Haliclona* species (Demospongiae, Haplosclerida) in the Austin and McDaniel collections at the Royal British Columbia Museum (RBCM) are identified only to genus or genus and species. The collections are representative of over 40 years of sampling principally by the late Dr. William C. Austin and one of us (Neil McDaniel) through SCUBA diving on the west coast of British Columbia and specimens provided by others to Dr. Austin. We have selected representative *Haliclona* species in the collections for detailed examination and placement in subgenera and species (where species were not identified). *Haliclona* is recognized to have several subgenera, thus identification of specimens to genus and species is incomplete. Our study updates this status for the species examined.

**Methods:** Methods of collection included intertidal scrapings or removal of non-encrusting specimens usually accompanied by in-situ photos, similar methods at SCUBA diving depths (subtidal to 35 m) and from other dredging, trawling and biological sampling activities.

**Results:** We describe eleven new *Haliclona* (Demospongeae Haplosclerida Chalinadae) species and a range extension for *Haliclona* (*Flagellia*) *edaphus* de Laubenfels, 1930 for shallow waters of Southwestern British Columbia, Canada. New species include *Haliclona* (*Gellius*) *hartmani* **n. sp.**, *Haliclona* (*Gellius*) *shishalhensis* **n. sp.**, *Haliclona* (*Reniera*) *gesteta* **n. sp.**, *Haliclona* (*Rhizoniera*) *aborescens* **n. sp.**, *Haliclona* (*Rhizoniera*) *blanca* **n. sp.**, *Haliclona* (*Rhizoniera*) *boothensis* **n. sp.,** *Haliclona* (*Rhizoniera*) *filix* **n. sp.**, *Haliclona* (*Rhizoniera*) *kunechina* **n.sp.,** *Haliclona* (*Rhizoniera*) *meandrina* **n. sp.,** *Haliclona* (*Rhizoniera*) *penelakuta* **n. sp.**, and *Haliclona* (*Rhizoniera*) *vulcana* **n. sp.** We also redescribe *Haliclona mollis* (Lambe, 1893 [1894]) and propose placing it in the subgenus *Haliclona*. Except for Lambe’s syntype slides of *Haliclona mollis* which are deposited at the Canadian Museum of Nature, Ottawa, Canada, all holotypes and voucher specimens of species described are deposited at RBCM.

## Introduction

In this report we describe eleven new *Haliclona* (Demospongeae Haplosclerida) species in the Royal British Columbia Museum (RBCM) collected over a 40-year period by William C. Austin and Neil McDaniel. Re-examination of *Haliclona* (*Flagellia*) *edaphus* de Laubenfels, 1930 specimens in the collection confirms their presence in British Columbia from previously reported Washington State, USA (de Laubenfels 1961). We also redescribe *Haliclona mollis* (Lambe, 1893 [1894]) from syntype slides loaned to RBCM by the Canadian Museum of Nature (CMN). The new species are all from littoral and shallow water locations in southwest British Columbia (Figure 1) and many fairly commonly encountered. To date most BC *Haliclona* have only been identified to genus or genus and species. De Weerdt (1989) proposed a major revision of the family Chalinidae that included erecting six subgenera in *Haliclona*; Van Soest (2017) added *Flagellia* to de Weerdt’s *Gellius, Halichoclona, Haliclona, Reniera, Rhizoniera* and *Soestella*. We selected *Haliclona* to provide some insight into the diversity of this sponge below the genus level in British Columbia waters.

**Figure 1.**
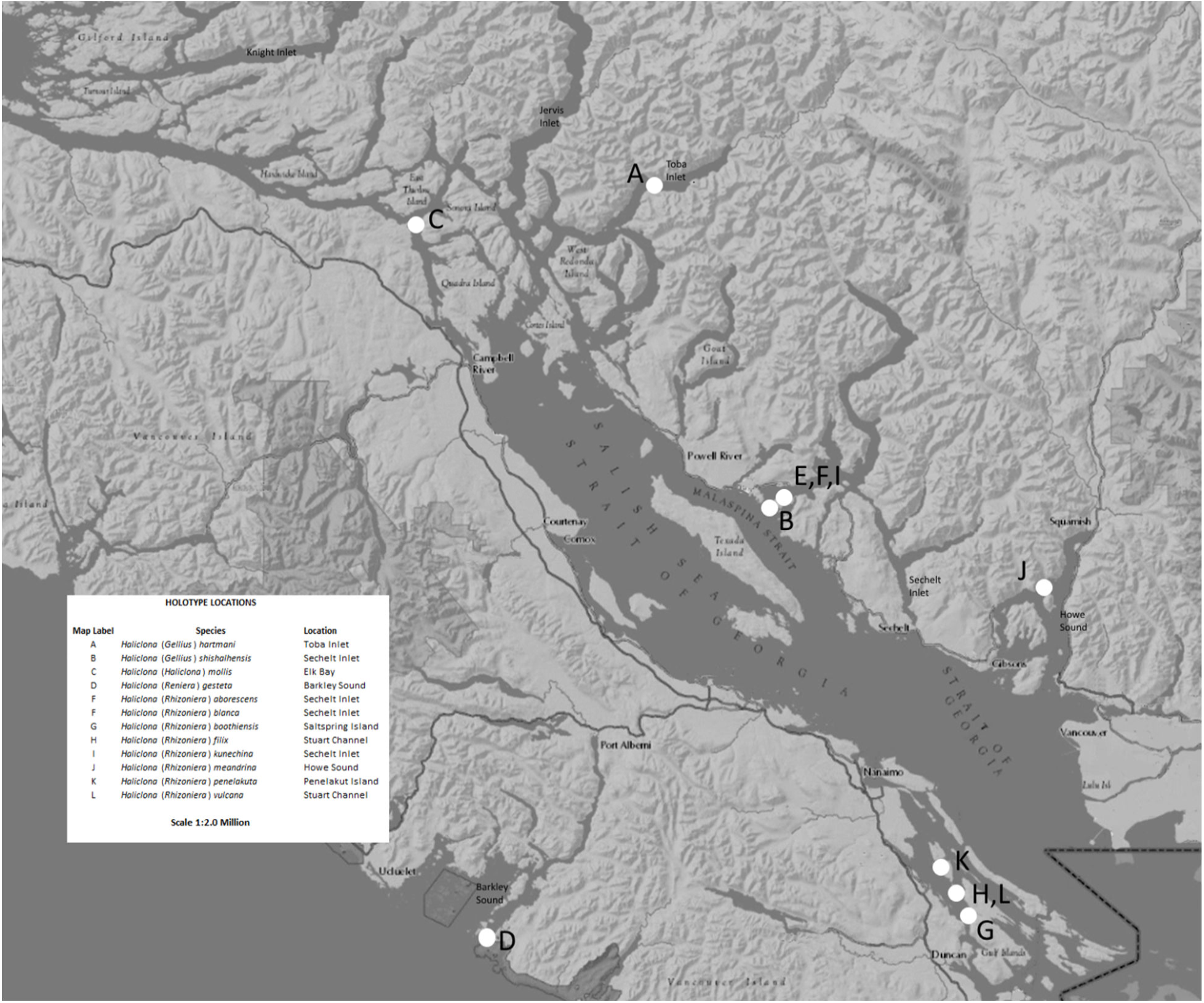
Holotype Location Map (Adapted from a Government of BC map, printed with permission).

*Haliclona* is representated by four subgenera and unclassified to subgenus in BC waters where collections have been made. Previously recorded *Haliclona* species specifically in the southwest BC marine area (based on unpublished species lists compiled by the late Dr. W.C. Austin) include *Haliclonia* (*Flagellia*) *edaphus* (de Laubenfels, 1930), *Haliclona* (*Flagellia*) *porosa* Fristedt 1887, *Haliclona* (*Gellius*) species cf. *emiltopsenti* Van Soest & Hooper, 2020 [formerly *H*. (*G*.) *foraminosa* (Topsent, 1904)], *Haliclona mollis* (Lambe, 1893) and *Haliclona* species cf. *mollis* (Lambe, 1893).

This report adds two new *H*. (*Gellius*) species, one new species of *H*. (*Reniera*) and eight new *H*. (*Rhizoniera*) species to *Haliclona* species previously recorded for British Columbia. The new species include: *Haliclona* (*Gellius*) *hartmani* **n. sp.**, *Haliclona* (*Gellius*) *shishalhensis* **n. sp.**, *Haliclona* (*Reniera*) *gesteta* **n. sp.**, *Haliclona* (*Rhizoniera*) *aborescens* **n. sp.**, *Haliclona* (*Rhizoniera*) *blanca* **n. sp.**, *Haliclona* (*Rhizoniera*) *boothiensis* **n. sp.**, *Haliclona* (*Rhizoniera*) *filix* **n.sp.**, *Haliclona* (*Rhizoniera*) *kunechina* **n.sp.**, *Haliclona* (*Rhizoniera*) *meandrina* **n. sp.,** *Haliclona* (*Rhizoniera*) *penelakuta* **n. sp.** and *Haliclona* (*Rhizoniera*) *vulcana* **n. sp.**

## Material and Methods

Methods of collection included intertidal scrapings or removal of non-encrusting specimens usually accompanied by in-situ photos, similar methods at SCUBA diving depths (subtidal to 35 m) and from other dredging, trawling and biological sampling activities. Skeletal thick sections and tissue-free spicule slides were prepared as described in Austin, et al. (2014). Spicule micrographs were made with a compound light microscope and camera at RBCM. Spicule dimensions are in microns (µm) as minimum (average) maximum. Number measured = 50 unless indicated. In spicule dimension tables holotypes are listed first and stations bolded. Scale bars on in-situ figures are approximate. Abbreviations: BO Bruce Ott; CMN Canadian Musuem of Nature, Ottawa, Canada; KML Khoyatan Marine Laboratory; NM Neil McDaniel; PBS Pacific Biological Station, Nanaimo, BC.; PEI Pacific Environment Institute of Fisheries & Oceans Canada; RBCM Royal British Columbia Museum; RH Rick Harbo; VT Verena Tunnicliffe.

## Systematics

Sponge classification follows that of Morrow & Cárdenas (2015) and specifically for Haplosclerida that of de Weerdt (2002 [2004]) and Van Soest (2017).

Demospongia Sollas, 1885

Haplosclerida Topsent, 1928

Chalinidae Gray, 1867

*Haliclona* Grant, 1841

*Haliclona* (*Flagellia*) Van Soest, 2017

## Synonymy

For world synonymy and distribution see Van Soest (2017).

### *Haliclona* (*Flagellia*) *edaphus* de Laubenfels, 1930 (Figure 2)

*Haliclona* (*Flagellia*) *edaphus* (as *Sigmadocia edaphus*) was reported in a web-based database by the late Dr. W.C. Austin. Portions of Austin’s database are listed in an undated University of British Columbia on-line PDF and noted by Van Soest, 2017, p.7. We have re-examined the Austin specimens in the RBCM collections and provide data to support a range extension of the species to British Columbia. We include a brief description and spicule dimensions.

**Figure 2.**
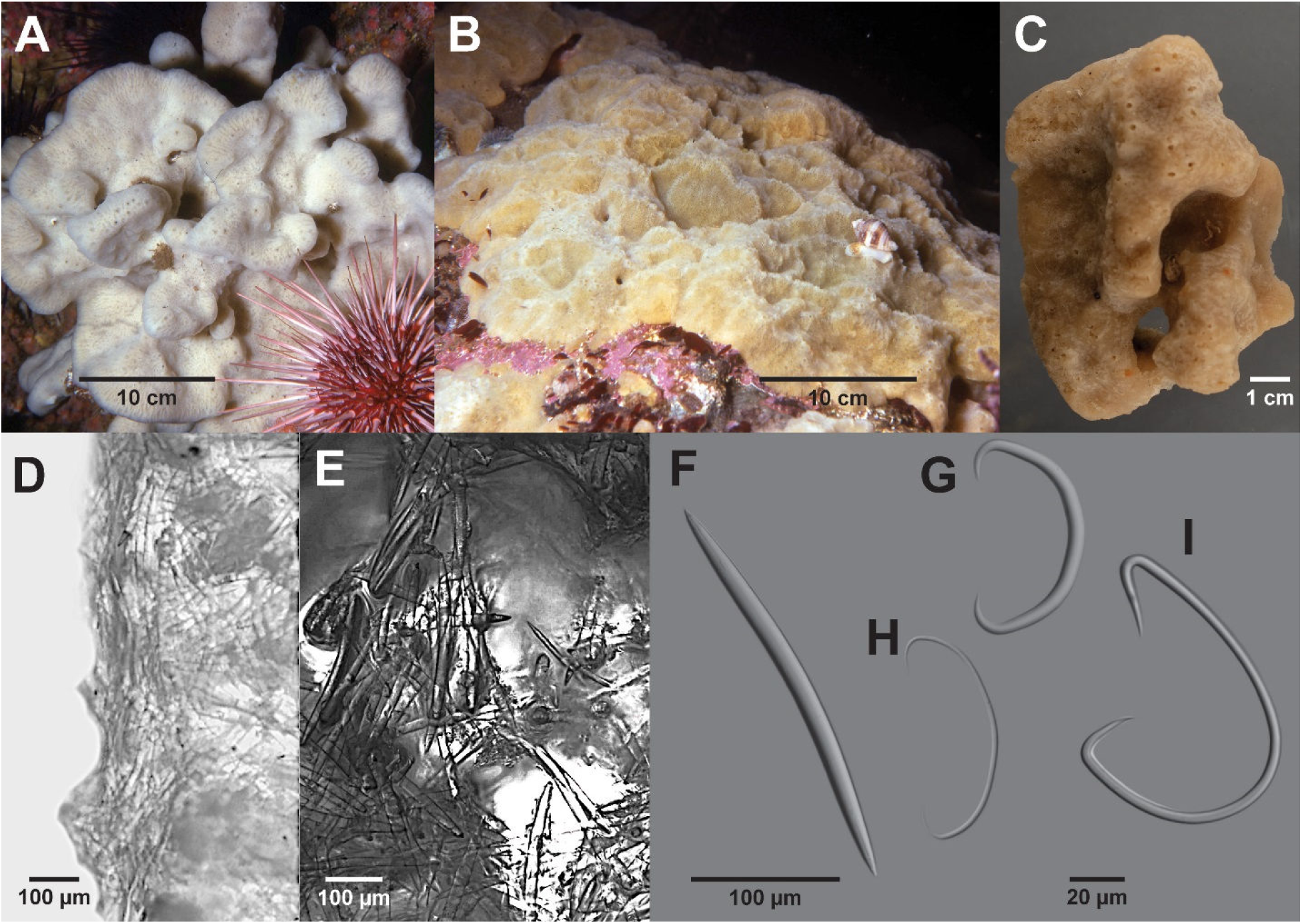
H*a*liclona (*Flagellia*) *edaphus*. A in-situ high energy current swept area. B in-situ high energy surf swept area. C whole preserved specimen. D ectosome and upper choanosome, oblique view. E choanosome. F oxea. G thick Normal sigma. H immature Normal sigma. I flagellosigma. A, B photographs only, no voucher specimens. C, RBCM 018-00262-001, D–I, RBCM 018-00262-002.

## Material Examined

RBCM 018-00262-001, station KML 85/73 off Whitlestone Point, Barkley Sound, 48° 49.28’ N / 125° 11.06’ W, 10–13 m depth, 5 May 1973, collector W.C. Austin, 1 specimen. RBCM 018-00262-002, same location and station, 1 specimen. RBCM 018-00278-001, station KML 90A/73, Barkley Sound, cave, 48° 48.8’ N / 125° 10.6’ W, +3 m, 5 Aug 1973, collector W.C. Austin, 1 specimen. RBCM 018-00381-001, station KML 240/70, Barkley Sound, 48° 48.9’ N / 125° 10.7’ W, 12 m depth, 21 Sep 1970, collector W.C. Austin, 1 specimen. RBCM 018-00382-001, station KML 89/77, James Island, near Sidney, 48°36.18’ N / 123°21.00’ W (approximate), depth not recorded, 27 Mar 1977, collector W.C. Austin, 1 specimen. Station PBS 1960, Graham Island, Haida Gwaii, BC, 54° 06.09’ N 132° 26.24’ W, 66 m depth, 24 Sep 1960, collector PBS, Nanaimo, BC, 1 non voucher specimen.

## Description

RBCM 018-00262-001, station KML 85/73 is representative of the Austin specimens and the best preserved. Representative living specimens were photographed by one of us (N.M.) are hard, spreading thickly encrusting and a shade of white.

**External** (Figure 2A) Described from an alcohol-preserved specimen. Massive, irregular shape, 88 mm L x 55 mm W x 60 mm H. Oscula on low conules, 2 mm diameter. Colour in alcohol light brown. Consistency firm, preserved specimen easily torn.

**Skeleton** Ectosome (Figure 2D) Discontinuous layer of oxeas parallel to surface with oxeas penetrating at various angles randomly along the surface. Ectosome 50 µm thick. Oxeas penetrate up to 200 µm. Choanosome (Figure 2E) Vague multispicular reticulation of oxeas around widely-spaced cavities a few hundred microns in diameter.

**Spicules** Spicules are oxeas including immature stages (abundant), Normal sigmas including immature stages (common), and flagellosigmas (uncommon). Stylote forms of oxeas occur occasionally. Oxeas (Figure 2F) are curved, have a cylindrical shaft and acerate apices slightly less than one fifth the total oxea length. Immature oxeas are about the same length as fully developed ones but thinner. Normal sigmas (Figure 2G) have a generally uniform arc with a 0.06 width to chord length ratio and sharp, strongly recurved apices. Immature Normal sigmas, while also C-shaped, have a more variable arc (typical form shown in Figure 2H), a width to chord length ratio of approximately 0.03; apices are similar to thick Normal sigmas. Flagellosigmas (Figure 2I) are typical for the subgenus (Van Soest 2017). Spicule dimensions of BC specimens are listed in Table 1.

**Table 1.** *Haliclona* (*Flagellia*) *edaphus* BC Specimens Spicule Dimensions (µm)

| Specimen | Oxeas | Normal sigmas<br>fully developed | Normal sigmas<br>immature | Flagellosigmas |  |  |  |
| --- | --- | --- | --- | --- | --- | --- | --- |
|  |  |  |  | Long Length | Short Length | Width | Thickness |
| KML 85/73<br>RBCM 018-00262-002 | 242 (284) 315 x<br>11.7 (18.1) 21.0 | 33.8 (52.1)<br>68.9 | 44.2 (70.0)<br>96.2 | 70.2 (95.9)<br>107<sup>a</sup> | 36.4 (63.4)<br>72.8 | 46.8 (73.1)<br>93.6 | 2.6 (4.0) 7.3 |
| KML 90A/73<br>RBCM 018-00278-001 | 245 (285) 315 x<br>8.8 (16.9) 26.3 | 15.6 (46.5)<br>65.0 | 9.0 (64.7) 125 | 78.0 (89.8)<br>104<sup>a</sup> | 52.0 (60.5)<br>70.2 | 54.6 (63.3)<br>78.0 | 2.3 (2.7) 3.6 |
| KML 240/70<br>RBCM 018-00381-00 | 175 (273) 315 x<br>9.6 (17.1) 21.9 | 23.4 (53.6)<br>70.2 | 39.0 (68.5) 109 | 54.0 (88.0)<br>109<sup>b</sup> | 36.4 (62.6)<br>80.6 | 52.0 (69.1)<br>93.6 | 2.6 (4.2) 5.2 |
| KML 89/77<br>RBCM 018-00382-001 | 221 (300) 336 x<br>10.4 (20.3) 23.4 | 39.0 (55.8)<br>83.2 | 33.8 (67.8)<br>88.4 (n=40) | 57.2 (81.9)<br>101<sup>c</sup> | 49.4 (55.9)<br>65.0 | 59.8 (71.5)<br>88.4 | 4.7 (5.2) 5.5 |
| PBS 1960<br>non voucher<br>specimen | 263 (297) 357 x<br>13.0 (17.2) 19.5 | 44.2 (72.7)<br>91.0 | 23.4 (38.5)<br>49.4 | 75.4 (92.2) 112 | 39.0 (61.3)<br>80.6 | 41.6 (74.1) 101 | 1.3 (3.0) 5.2 |

<sup>a</sup> flagellosigmas n=15 <sup>b</sup> flagellosigmas n=14 <sup>c</sup> flagellosigmas n=6 <sup>d</sup> flagellosigmas n=17

## Distribution

British Columbia (BC), west coast of Vancouver Island, intertidal to Graham Island, Haida Gwaii, BC 66 m. The species in British Columbia is found on both inner and outer coasts.

## Remarks

BC preserved specimens fit fairly closely the type description (de Laubenfels, 1932 as amended by Lee, et al. 2007 and Van Soest 2017) with the following exceptions:

- oscula are up to 3 mm diameter vs. about 1 mm for California specimens, possibly a preservation artifact of the BC specimens;
- oxeas are larger (to 389 x 26 µm vs. 300 x15 µm of California specimens) possibly explained by higher silica concentration in the BC sea water (Austin, et al. 2014, p. 8);

### Haliclona (Gellius) Gray, 1867

#### World Distribution

There are 78 *Haliclona* (*Gellius*) subspecies world-wide (de Voogd, et al. 2024). Nine species of *H.* (*Gellius*) are reported for the North Pacific and three for the Northeast Pacific; two *Haliclona* no subgenus species with at least oxeas and sigmas are also reported for the Northwest Pacific (Japan) (Table 2).

**Table 2.** North Pacific *Haliclona* (*Gellius*) and *Haliclona* No Subgenus Species with Oxeas and Sigmas.

| Species | Location | Depth (m) | External | Skeleton | Spicules |
| --- | --- | --- | --- | --- | --- |
| *Haliclona* (*Gellius*) cf. *cymiformis* (Esper, 1806). | Barkley Sound | Littoral | Erect, branching, encrusting. Surface smooth. Oscula flush, 0.8–1.5 mm. Colour in life dark green. | Unispicular isodictyal reticulation in ectosome; may be broken. Choanosome: unordered except 5–6 spicule short tracts. No spongin. Sigmas throughout sponge. | Oxeas: curved, long, sharp points, 128–160 x 2–7.5 µm. Sigmas: C, 16–20 µm. |
| *Haliclona* (*Gellius*) *emiletopsentii* Van Soest & Hooper, 2020 [formerly *Haliclona* (*Gellius*) *foraminosa* (Topsent, 1904)] | Barkley Sound | 10<br>Topsent<br>200 | Thickly encrusting; surface smooth. Colour in life yellow-grey. | Ectosome: thin membrane. Choanosome: loose unispicular network w/ small amount of spongin at intersects. | Thick oxeas: slightly curved, sharp points 435 x 16 µm. Slender oxeas: 435 x 3–5 µm. |
| *Haliclona* (*Gellius*) *laubenfelsi* Van Soest & Hooper, 2020 [formerly *Haliclona* (*Gellius*) *violacea* (de Laubenfels, 1950)] | Hawaii | 1 | Encrusting; oscula raised, sometimes on fistulae, 3 mm diameter. Ostia abundant, 30 µm diameter. Smooth surface. Translucent dermis over subdermal cavities. Colour in life violet. | Not recorded. | Oxeas: 120–140 x 4–7 µm. Toxas: 60 µm. |
| *Haliclona* (*Gellius*) *microxea* (Li, 1986) | China, Yulin | No data | The sponge is encrusting, attached on the shell of bivalves. No colour in life. | The skeleton consists of rather close-meshed reticulation of unispicule, the meshes are triangular or quadrilateral, the sides 100–150 µm long. | Oxea: two kinds, slightly curved, 154–196 x 3–7 µm; straight and finely echinated, slender raphides, 92–103 x 2–3 µm. Toxas: 70–86 x 2–3 µm. Sigmas: 14–22 x 1–2 µm. Microxeas: echinated, 33–36 x 2 µm. |
| *Haliclona (Gellius) primitiva* (Lundbeck, 1902) of Koltun (1958) | Southern Kuril Islands, western coast of S. Sakhalin Island, Sea of Okhotsk, Bering Sea | 27–40 | From Lundbeck: Encrusting, May be tubular. Dried yellowish. | From Lundbeck: Regular network, partly unispicular, but also polyspicular tracts, especially running in the direction towards the surface. The skeletal meshes are more or less rectangular. Especially the tracts running towards the surface (the primary ones) are distinct, while the tracts running vertically are less conspicuous. In the nodes of the skeleton the spicules are united by a distinct and rather copious mass of spongin. | Koltun Oxeas: curved, sharp points, 140–184 x 6–13 µm. Sigmas: C, 28–170 x 1–4 µm. |
| *Haliclona (Gellius) toxia* (Topsent, 1897) of Li (1986) | China, Beibu Gulf | No data | The sponge is irregularly massive, with some oscular tubes on the surface, the oscula opening several millimeters under the tip of the tube, 3–5 mm in diameter. No colour in life. | The dermal skeleton consists only of scattered oxea placed tangentially, the choanosome is a very close and pretty uniform reticulation of single oxeas. | Oxeas: slender, slightly curved, 175–180 x 7–9 µm. Toxas: two sizes, 28–35 µm. 65–115 x 3–5 µm. |
| *Haliclona (Gellius) varia* (Bowerbank, 1875) of Li (1986) | Hong Kong | No data | The sponge is attached below by an encrusting base, and forms an erect, leaf like plate branched, 1–2 mm thick. Oscula not visible. | The dermal skeleton consists of scattered area, typical of Halichondria, The choanosome is a loose, rather irregular reticulation. | Oxeas: slightly curved, end of some spicules variously shape, pointed, 212–215 x 5–11 µm. Sigmas: C, 22–31 x 2 µm. |
| *Haliclona (Gellius) vladimirkoltuni* Van Soest & Hooper, 2020 [formerly *H. (G.) digitata* (Koltun, 1958)] | Sea of Okhotsk | 285–287 | The sponge is elongated, thin-leaf-shaped, rolled up and fused in such a way that it forms an irregularly shaped body, hollow inside and narrowed toward the base with hollow finger-like outgrowths. Sponge up to 7 cm high. Color in life light yellow. | Skeleton consists of branched tracts cross-connected by individual spicules. | Oxeas: slightly curved, moderately short, sharp points, 320–370 x 15–18 µm. Sigmas: C, 18–23 x 1–2 µm. |
| *Haliclona (Gellius)* cf. sp. of Hartman, 1975 | Toba Inlet, Horseshoe Bay<br>Hartman: central Cal,<br>intertidal/subtidal | HB: 50 | From Lee, et al. 2007<br>Encrusting to 12 mm thick.<br>Colour in life rose lavender. | From Lee, et al. 2007<br>Ectosome: some oxeads at<br>surface of dermal layer; others,<br>stand on end and pierce the<br>surface. Sigmas and spongin<br>abundant in surface layer.<br>Subdermal cavities present.<br>Vertical unispicular and<br>paucispicular tracts descend<br>into the choanosome with<br>horizontal and diagonal cross-<br>spicules. Deeper in the<br>choanosome a greater number<br>of confused spicules occur, but<br>a basic isodictyal pattern<br>remains. | From Lee, et al. 2007<br>Oxeas: means, 170–190 x 7.0–<br>9.0 µm. Sigmas: C-shaped 24–<br>36 µm. |
| *Haliclona liber* (Hoshino, 1981) | Japan | 12–13 | Very thin encrusting. Surface<br>smooth, even. Colour in life<br>grey. | No ectosome. Choanosome<br>loose, irregular reticulation.<br>Primary tracts 30–50 µm thick.<br>Secondary tracts 10–20 µm<br>thick. Sigmas near tracts or in<br>flesh. | Oxeas: slightly arched or bent<br>in middle, tapering to apices,<br>150–175 x 6–8 µm. Sigmas: C,<br>thin, 15 µm. |
| *Haliclona uwaensis*<br>(Hoshino, 1981) | Japan | Subtidal | Very thin encrusting. Oscula<br>not visible. Ivory buff dry. | No ectosome. Choanosome<br>irregular coarse network of<br>spiculo-fibre from 1–10 rows<br>of spicules, 30 µm diameter.<br>Numerous toxas and sigmas<br>scattered in flesh. | Oxeas: slightly arched, hastate,<br>tapering to apices, 150–185 x<br>6–10 µm. Toxas: thin, 28 µm.<br>Sigmas: C, 10–22 µm. |

#### *Haliclona (Gellius)* hartmani n. sp. (Figure 3)

Zoobank yyyy

**Figure 3.**
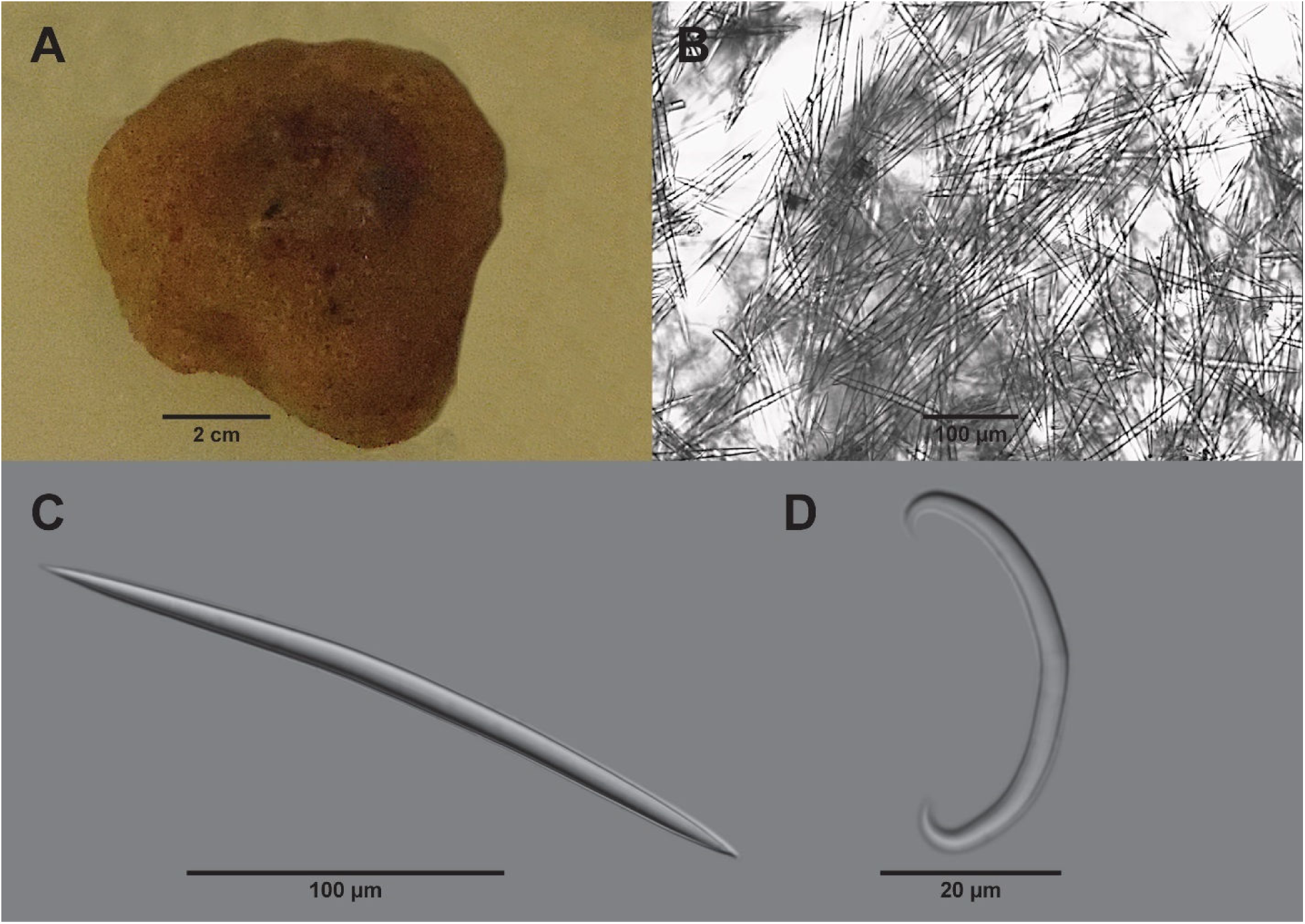
H*a*liclona (*Gellius*) *hartmani* n. sp. A-D: Holotype, RBCM 014-00223-008, A Sponge top view. B Skeleton cross section. C Oxea. D Normal sigma.

**Diagnosis** Encrusting, porous sponge with no ectosome and a halichondrid choanosome.

**Etymology** The sponge is named after Dr. Willard Hartman, who identified a similar sponge to genus from central California.

## Material Examined

Holotype RBCM 014-00223-008, station KML 241/82, Toba Inlet, BC, S side approximately 10 km from the entrance, 50° 24.671’ N / 124° 30.531’ W 15 m depth, 5 Dec 1982, collector W.C.

Austin, 1 specimen.

## Description

**External** Described from an alcohol-preserved specimen. Sponge irregularly massive, 3 cm on a side. Oscula scattered, slightly raised on shallow conules 2–3 mm diameter. Surface rough to touch. Colour light brown in alcohol. Consistency firm, not easily compressed (Figure 3A).

**Skeleton** No specialized ectosome. Skeleton a confused arrangement of oxeas with vague multispicular tracts. Sigmas scattered throughout (Figure 3B).

**Spicules** Spicules include oxeas and Normal sigmas. Oxeas are usually curved with moderately long slightly mucronate apices, occasionally straight, 192 (225) 255 x 7.8 (10.1) 13.0 µm, abundant (Figure 3C). Immature oxeas common. Sigmas C-shape, 33.8 (45.1) 62.4 µm, abundant (Figure 2D).

**Distribution** Found only at the type locality, approximately 10 km from the entrance to Toba Inlet, BC in 15 m of water; may be conspecific with Hartman’s *Haliclona* (*Gellius*) species from central California.

## Remarks

The sponge was originally labeled by Austin in 1982 as *Sigmadocia* species of Hartman 1975 (based on the station label with the specimen). Based on the description of Hartman’s sponge by Lee, et al. (2007, p. 228), the BC sponge has a similar habitus (except Hartman’s sponge is rose-lavender and our BC specimen is white), skeletal structure and spicule types, but larger spicules (BC oxeas to 255 µm vs California oxeas to 190 µm; BC sigmas to 62 µm vs California sigmas to 36 µm). There are no published records of Hartman’s sponge from Oregon or Washington.

Comparisons with the brief descriptions in Table 2, none of the other listed species is sufficiently similar to *H.* (*G.*) *hartmani* **n. sp.** to be considered conspecific.

*Haliclona (Gellius) shishalhensis* n. sp. (Figure 4)

**Figure 4.**
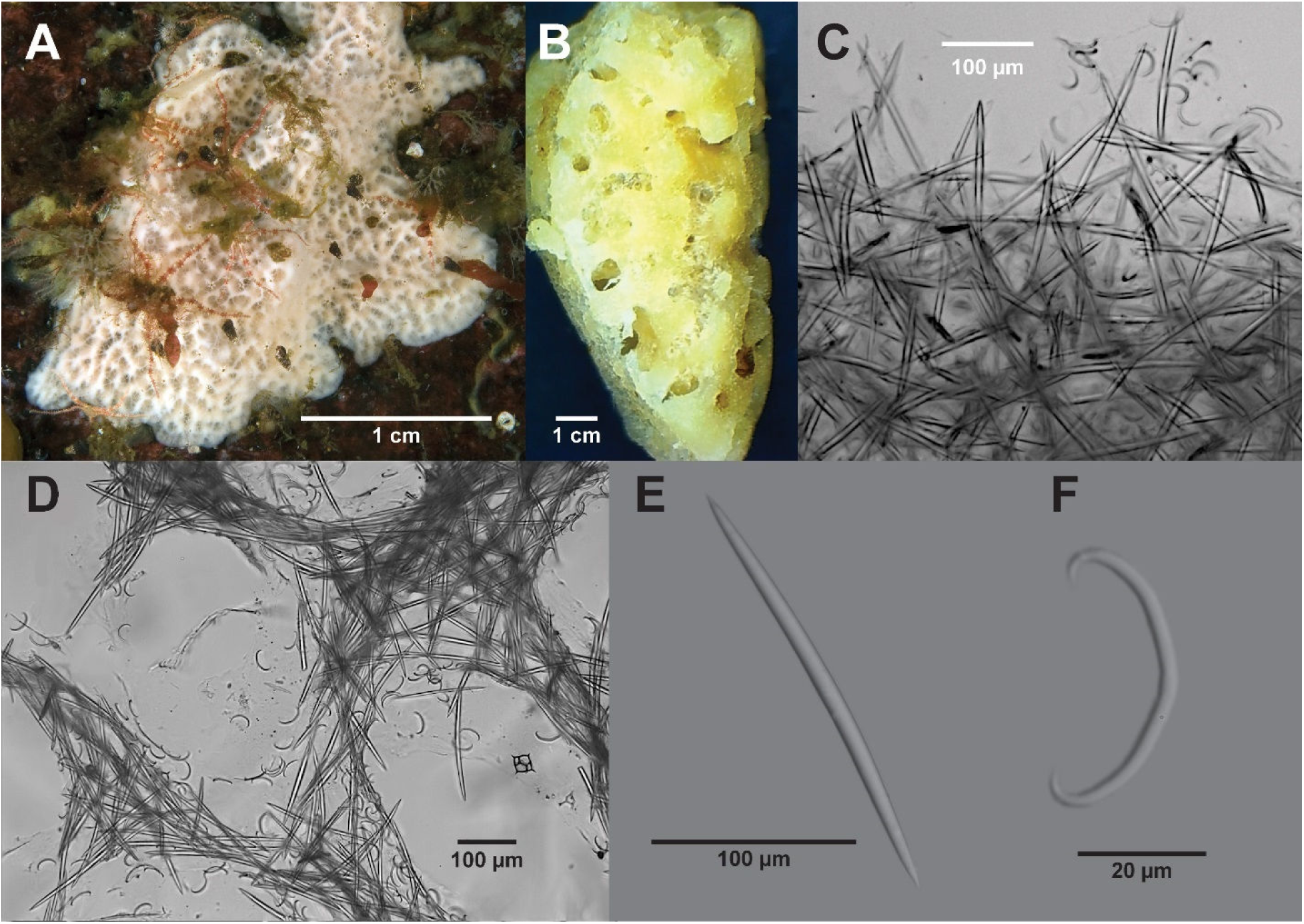
H*a*liclona (*Gellius*) *shishalhensis* **n. sp.** A-F Holotype, RBCM 018-00383-001. A Holotype in-situ. B Cross section showing cortex-like surface with large lacunae beneath. C Skeleton cross section at surface. D Skeleton cross section in open part of choanosome. E Oxea. F Normal sigma.

Zoobank yyyy

**Diagnosis** Very open porous structure with a ridgid unseparable ectosome and a ridged surface. No ectosome, choanosome a vague anisotropic reticulation around large, numerous aquiferous canals.

**Etymology** The species name means from Shíshálh (English: Sechelt) home of the Shíshálh People, the location of the specimens described herein. Per the International Code of Zoologic Nomenclature (4^th^ Ed., 1999, Article 27) diacritical marks have been dropped from the species name.

**Material Examined** Holotype RBCM 018-00383-001, station NM 341, Sakinaw Rock, Sechelt Inlet, BC, 49° 33.947’ N / 123° 48.222’ W, 15 m, 26 July 2016, collector N. McDaniel, 1 specimen. Paratype RBCM 018-00152-006, station NM 240, Nine Mile Point, Sechelt Inlet, BC, 49° 36.293’ N / 123° 47.139’ W, 23 m depth, 12 May 2011, collector N. McDaniel, 1 specimen.

## Description

**External** (Figure 4A) Sponge thickly encrusting 4 x 2 x 1.5 cm. Very porous, open structure; ectosome 1 mm thick, fairly rigid, not detachable. Pores 100–300 µm diameter (preserved). Area around pores slightly raised resulting in a network of ridges. Oscula are not evident (seprable from pores). In several places the sponge is completely hollow from the surface to the substrate. Surface microhispid; spicules project 100 µm. White in life. Fairly easily torn.

**Skeleton** There is no specialized ectosome structure but macroscopically a more dense cortex-like layer is formed in the upper 1 mm. Subdermal lacunae 0.3 to 1.5 mm wide connect to surface pores (Figure 4B). Single, or one to two oxeas on a side form an irregular anisotropic reticulation with single oxeas slightly penetrating the surface up to 100 µm (Figure 4C). The choanosome consists of short, multispicular tracts or compressed anisotropic reticulations around large aquiferous canals (Figure 4D). In areas away from aquiferous canals an anisotropic reticulation or occasional short multispicular tracts are formed. Aquiferous canals deeper in the sponge average 400 µm diameter. Sigmas are located throughout the sponge. Spongin at nodes scarce to absent.

**Spicules** Megascleres are oxeas, gently curved with sharp apices, 177–226 x 6.5–13 µm (Figure 4E). Microscleres are C sigmas, uniformly curved or slightly bent at the centre, 16.9–33.8 µm (Figure 4F). Dimensions of the two specimens examined are listed in Table 3.

**Table 3:** *Haliclona* (*Gellius*) *shishalhensis* **n. sp.** Spicule Dimensions (µm)

| Station | Spicule | Length | Width |
| --- | --- | --- | --- |
| NM 341<br>RBCM 018-00383-001 | Oxeas | 182 (206) 226 | 6.5 (8.7) 13.0 |
|  | Sigmas | 16.9 (22.7) 33.8 |  |
| NM 240<br>RBCM 018-00152-006 | Oxeas | 177 (197) 216 | 7.8 (8.7) 10.4 |
|  | Sigmas | 20.3 (22.5) 27.0 |  |

**Distribution** Found at two locations in Sechelt Inlet, BC about 5 km apart and at 15 and 23 m depths.

## Remarks

Two *Haliclona* (*Gellius*) species were reported by W.C. Austin (unpublished northeast Pacific sponge list) for BC: a species similar to *H.* (*G.*) *foraminosa* Topsent, 1904 (now *H.* (*G.*) *emiletopsenti* Van Soest & Hooper, 2020) in Jervis Inlet and a species similar to a sponge from central California identified by Hartman (1975) as *Sigmadocia = Haliclona* (*Gellius*) per De Weerdt (2002 [2004]) at Horseshoe Bay (since lost) and Toba Inlet. Topsent’s sponge has two sizes of oxeas and no sigmas (see Table 2) and was from the Azores, 200 m depth. Based on the description of Hartman’s sponge provided by Lee et al. (2007, p. 228) Hartman’s sponge does not have a cortex-like ectosome.

### Haliclona (Haliclona) Grant, 1836

*Haliclona* (*Haliclona*) *mollis* n. sgen. (Lambe, 1893 [1894]) (Figures 5, 6)

**Figure 5.**
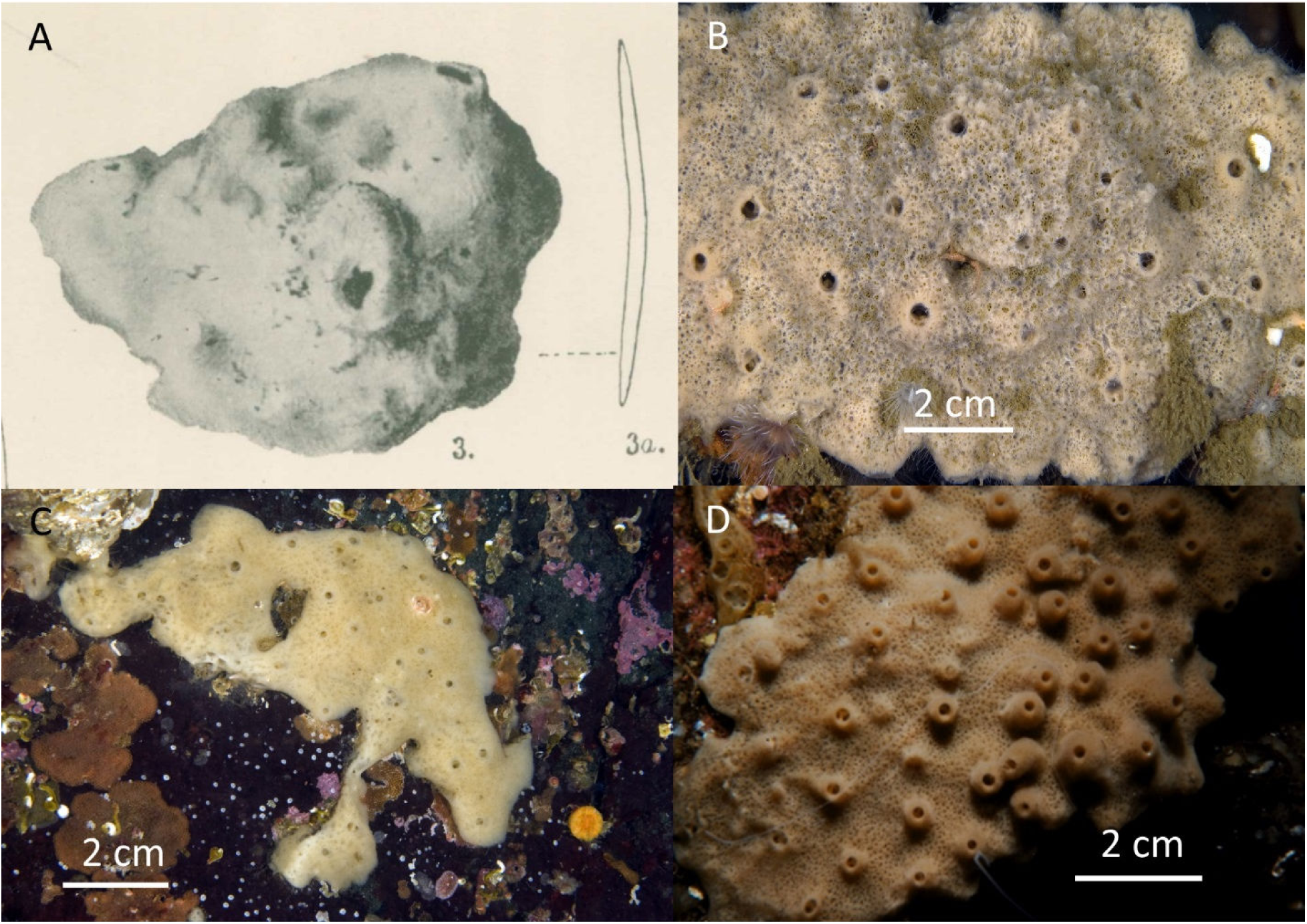
H*a*liclona (*Haliclona*) *mollis* A Lambe’s holotype, CMN Cat. No. CMNI 1900-2875. B–D In-situ representative BC specimens.

**Figure 6.**
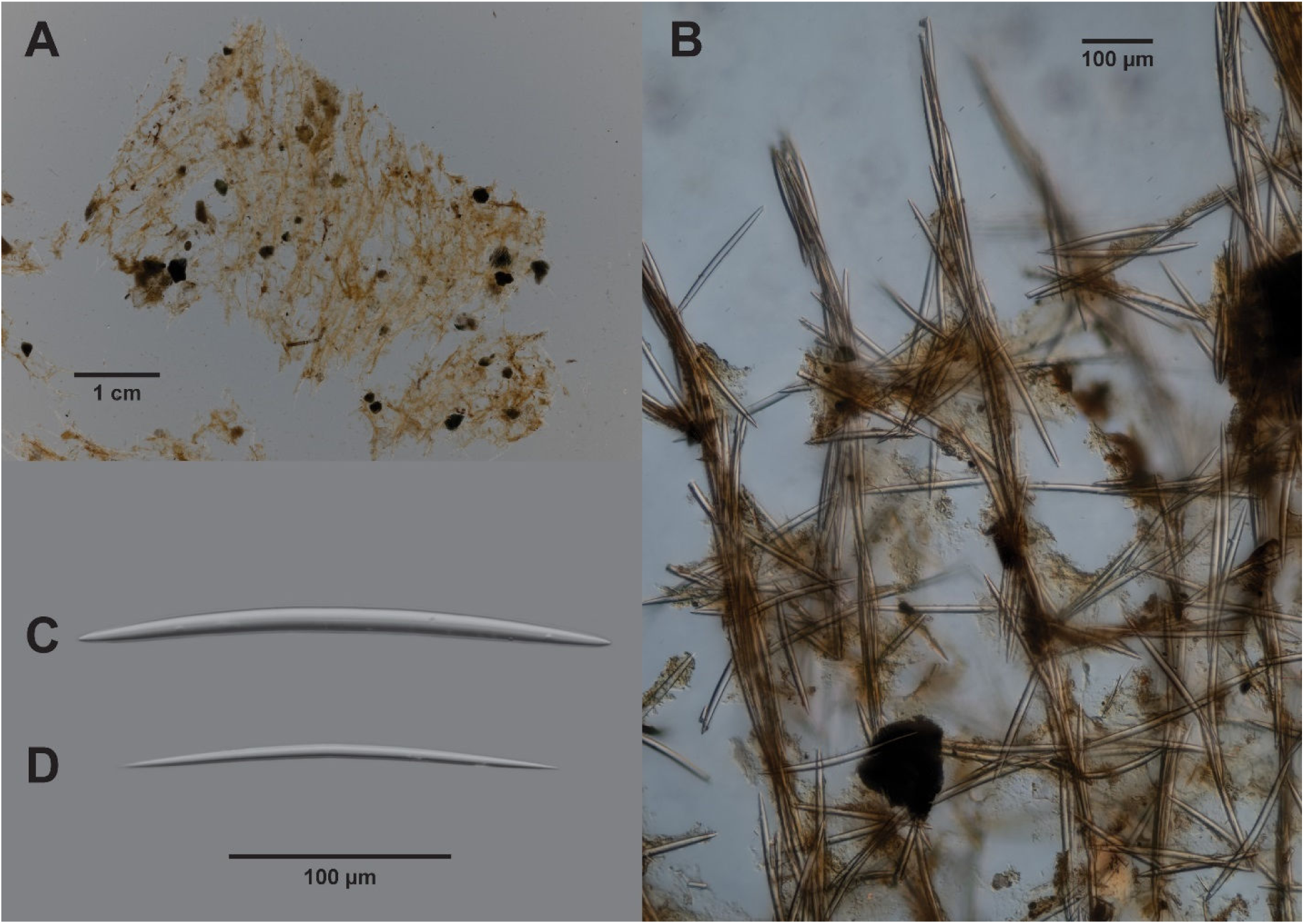
H*a*liclona (*Haliclona*) *mollis*. A–D, CMNI 1900-2875. A Skeleton cross section; surface upper right. B Skeleton close up near surface. C Oxea. D Immature oxea.

Zoobank yyyy

We propose the reclassification of Lambe’s *Haliclona mollis* (originally *Reniera mollis*) as *Haliclona* (*Haliclona*) *mollis* (Lambe, 1893 [1894]) based on a reexamination of Lambe’s slides of his large and small specimens provided courtesy of the Canadian Museum of Nature (CMN).

**Material Examined** Six syntype slides made by Lambe and labelled *Reniera mollis* (either larger or small specimen), Labels indicate Elk Bay, Dis. Passage [Discovery Passage], 20–25 fath[oms] [40–50 m], GMD [G.M. Dawson], 23^rd^ Jul/85 [23 July 1885]. CMN Cat. No. CMNI 1900-2875.

## Description

**External** (from Lambe’s paper, p. 26, pl. II, f. 3, 3a) *Sponge massive, sessile, growing in sublobate masses. Represented in the collection by two specimens, one 90 mm. long, 55 mm. high and 33 mm. thick, the other (Plate II, fig. 3) much smaller, 50 mm. long, 33 mm. broad and 30 mm. high. Colour in spirit, dull brownish-yellow. Texture soft and fragile. Surface uneven, hispid. Dermal membrane thin, aspiculous. Oscula large, prominent, attaining a diameter of 5 mm; in the larger specimen the oscula form an indistinct uniserial row along the sides, but in the smaller specimen they are irregularly disposed. Pores, appearing as circular or oval openings in the dermal membrane over large subdermal cavities. They are about 0 065 mm. in width and less than their width apart.* Lambe’s figure is reproduced here as Figure 5A. Habitus of specimens photographed in-situ in BC vary and are discussed below.

Lambe’s specimens were massive whereas BC specimens shown (and reviewed) are thickly encrusting, but otherwise identical to Lambe’s sponge. In thinner specimens (Figure 5C) oscula are not collared but approximately the same size. Colour in life varies from yellow to beige to light brown. Consistency live is soft, easily torn.

**Skeleton** (described from the syntype slides)

The skeleton consists of multispicular tracts running vertically from the base to the surface and projecting slightly beyond the surface 200–300 µm (Figure 6A). The aspiculous dermal membrane mentioned by Lambe is not visible in the slides. Tracts are composed of 3 to 7 spicules, crossed at regular intervals by single spicules (occasionally up to 3), forming a rectangular reticulation (Figure 6B). The distance between the principal vertical tracts is 127 to 297 µm (mean 208 µm) and the distance between horizontal cross spicules is 97 to 273 µm (mean 176 µm). Disposed among the tracts are numerous aquiferous canals, 71 (161) 238 x 95 (312) 714 µm, n=20.

**Spicules** (described from the syntype slides). Spicules are exclusively oxeas, curved with sharp hastate apices (Figure 6C). Immature oxeas are relatively abundant and differ only in being thinner and (usually) shorter (Figure 6D). Our measurements are consistent with Lambe’s as are the other specimens included as examples of Southwest BC specimens (Table 4).

**Table 4.** Comparison of Lambe’s and This Paper Measurements (µm)

| Measurement Source | Length | Width | Number |
| --- | --- | --- | --- |
| Lambe Original | 196–262 | 13 |  |
| Lambe Syntype CMNI 1900-2875 | 206 (242) 260 | 7 (11.1) 13 | 50 |
| Other BC *H. (H.) mollis* |  |  |  |
| PEI 04 | 208 (236) 265 | 5.2 (10.5) 13.3 | 50 |
| NM 337 | 190 (225) 252 | 7.8 (10.3) 12.5 | 50 |
| NM 371 | 221 (239) 263 | 10.4 (12.2) 13.0 | 50 |
| Miles 2010.10.25 | 200 (237) 273 | 10.4 (12.0) 13.3 | 50 |
| KML 110/75 | 179 (212) 239 | 9.9 (12.7) 15.6 | 50 |

## Distribution

Based on examined specimens: southern BC, +0.6 m to 50 m.

### Haliclona (Reniera) Schmidt, 1862

*Haliclona (Reniera) gesteta* n. sp. (Figure 7)

**Figure 7.**
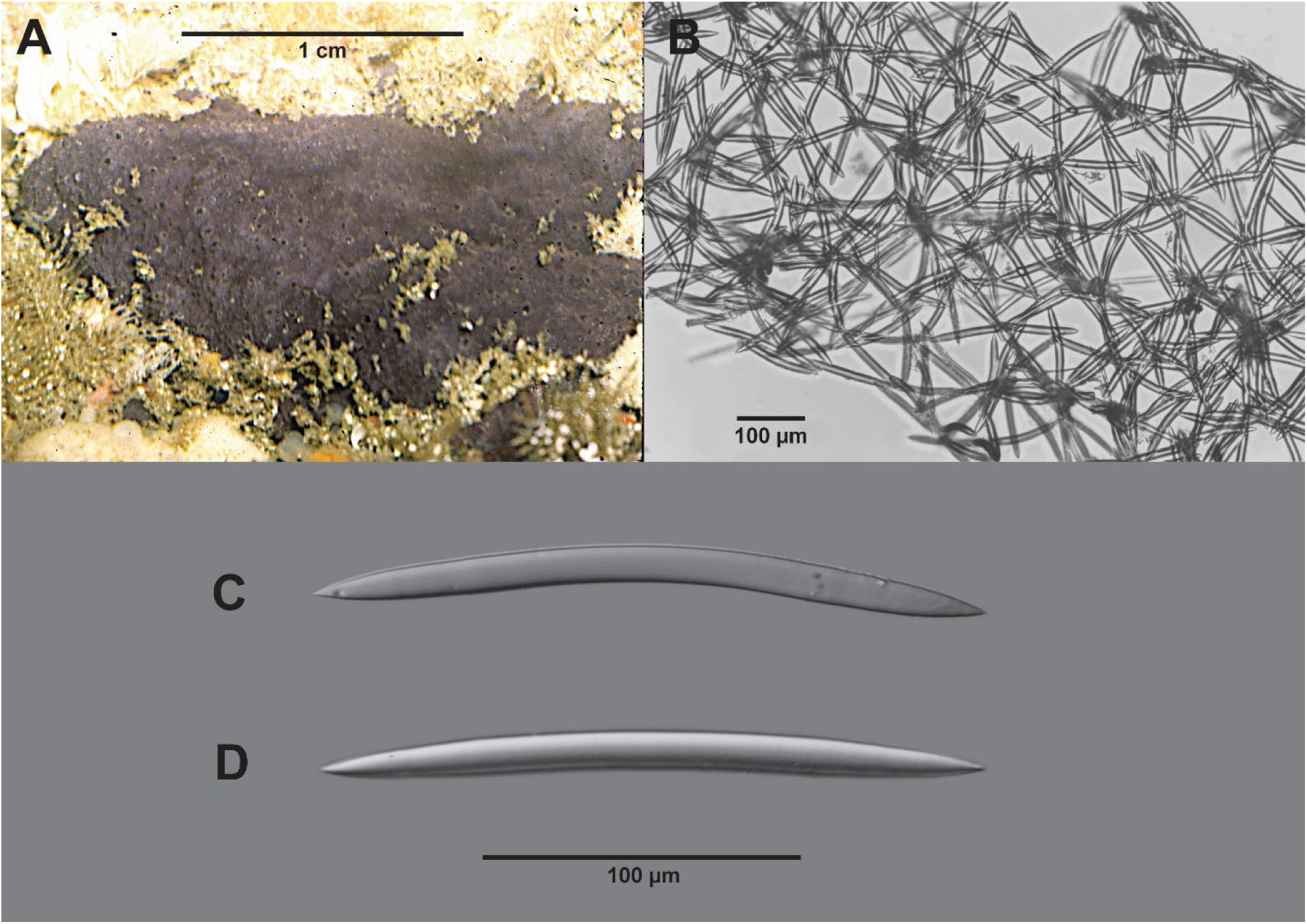
H*a*liclona (*Reniera*) *gesteta* **n. sp.** A–D, Holotype, RBCM 018-00272-001. A in-situ (photo W.C. Austin). B Skeleton cross section. C, D Oxeas.

Zoobank yyyy

**Etymology** The species name refers to its mauve colour.

**Diagnosis** Thin encrusting purple sponge with a micropapillate surface and small flush scattered oscula.

**Material Examined** Holotype RBCM 018-00272-001, station KML 80A/73, Execution Rock Cave, Barkley Sound, BC, 48° 49.9’ N / 125° 10.7’ W, littoral, 5 Aug 1973, collector W.C. Austin, 1 specimen.

## Description

**External** (Figure 7A) Thin encrusting, sponge 1.8 by 1.4 cm, base 0.8 mm thick. Micropapillae 2 mm high by 1 mm diameter, scattered. Oscula scattered about 3 mm diameter on live sponge. Base of sponge slightly rough. Colour mauve. Consistency soft, easily torn.

**Skeleton** The skeleton is typical renierid forming a subisotropic reticulation one spicule on a side with numerous loose spicules in the choanosome (Figure 7B). Spongin confined to some nodes. No evidence of spicule tracts. No specialized ectosome. Aquiferous canals 150 x 250 µm, oval, well-formed, not numerous [not figured].

**Spicules** (Figure 7C, D) Oxeas curved, hastate apices, few immature, rarely straight; rare styles, 140 (152) 164 x 10.4 (13.7) 15.9 µm.

## Distribution

Known from the type locality only, Execution Rock Cave, Barkley Sound, littoral.

## Remarks

The oxeas are typical of the subgenus as is the skeletal architecture. Austin (unpublished NE Pacific sponge list) identified two California species identified by Hartman (1975) as *Reniera* sp. A & B as occurring in BC. However, Lee et al. (2007) reclassified Hartman’s *Reniera* sponges as *Haliclona* (*Rhizoniera*) species. There are six accepted *H.* (*Reniera*) species described for the North Pacific (de Voogd, et al. 2024): *H*. (*R*.) *enormismacula* Hoshino, 1981 and *H*. (*R*.) *negro* (Tanita, 1965) from Japan; and *H*. (*R*.) *hongdoensis* Kang & Sim, 2007, *H*. (*R*.) *juckdoensis* Kim & Kang, 2020, *H*. (*R*.) *oceanus* Kim & Kang, 2020, and *H*. (*R*.) *sinyeoensis* Kang, Lee & Sim, 2013 from Korea. None are thin encrusting. *Haliclona* (*R*.) *sinyeoensis* is purple or pink but is cushion-shaped with oscula on chimneys.

There are 32 *Haliclona* no subgenera species reported for the North Pacific (de Voogd, et al. 2024). Ten are thin encrusting (listed in Table 5). *Haliclona liber* has sigmas, *H. uwaensis* has toxas and sigmas, *H. takaharui* has two sizes of oxeas. Skeletons of *H. hydroida*, *H. liber*, *H. offerospicula*, *H. tachibanaensis* and *H. takaharui* skeletons have spicule tracts. Of the remaining species, *H. densaspicula* is hard with visible oscula and colour is brown, *H. robustaspicula* has oscula up to 1 mm, cinammon pink colour and a skeleton with occasional tracts, *H. sataensis* is hard, old rose colour and a skeleton with occasional tracts, *H. scabrita* has a puntate smooth surface, light pinkish beige in colour, a skeleton with occasional tracts and oxeas smaller than *H. gesteta* **n. sp.** (130–170 x 8–9 µm).

**Table 5.** *Haliclona* No Subgenus North Pacific Species Similar to *H*. (*R*.) *gesteta* **n. sp.**

| *Haliclona* Species | Loc | Depth (m) | External | Skeleton | Spicules |
| --- | --- | --- | --- | --- | --- |
| *Haliclona densaspicula*<br>Hoshino, 1981 | Japan | Intertidal | Sponge 3x3x<0.5 cm. thick. Smooth, even. Oscules 0.2-0.5 cm diameter. Colour in life brown. Consistency hard, fragile. | No ectosome. Choanosome: isodictyal or subisodictyal reticulation. Numerous free oxeads. | Oxeads: hastate, straight or gently curved, 185–250 x 3–15 µm. |
| *Haliclona hydroida*<br>Tanita & Hoshino, 1989 | Japan | 20–25 | Encrusting, 1–2 mm. Surface smooth, punctiform, uneven. Oscula not visible. Colour (alcohol) ivory buff. Consistency compressible, not tough. | No ectosome. Choanosome regular network of primary and secondary tracts. Primary tracts 20 µm diameter, multispicular, ascending 100–200 µm apart. Secondary few rows of oxead between primaries at 500–100 µm. Primaries form surface brushes. | Oxeads: stout, hastate, 120–145 x 7–12 µm. |
| *Haliclona liber*<br>(Hoshino, 1981) | Japan | 12–13 | Very thin encrusting. Surface smooth, even. Colour grey. Consistency soft. | No ectosome. Choanosome: loose, irregular reticulation. Primary tracts 30–50 µm thick. Secondary tracts 10–20 µm thick. Sigmas near tracts or in flesh. | Oxeads: slightly arched or bent in middle, tapering to points, 150–175 x 6–8 µm. Sigmas: C, thin, 15 µm. |
| *Haliclona offerospicula*<br>Hoshino, 1981 | Japan | Intertidal | Small, irregular, thin. Smooth to touch. A few oscules, 2 mm diameter. Colour in life ivory buff. Consistency slightly compressible, fragile. | No ectosome. Choanosome: unispicular tracts 40–50 µm apart ascend to surface. Irregularly connected with oxeads. | Oxeads: straight to slightly curved with sharp points, 75–90 x 2–3 µm. |
| *Haliclona robustaspicula*<br>Hoshino, 1981 | Japan | Subtidal | Thin encrusting other sponges. Surface smooth or undulating. Oscules up to 1 mm loosely spaced. Colour in life cinammon pink. Consistency incompressible | No ectosome. Choanosome: irregular isodictyal reticulation w/ occasional irregular reticulation of bi and tri-spicule tracts. | Oxeads: hastate, gently curved or arched by bending twice at 1/3 spicule length, 265–295 x 11–13 µm. |
| *Haliclona sataensis*<br>Hoshino, 1981 | Japan | Subtidal | Irregular, thin, to 2 cm thick. Surface smooth, oscules not visible. Colour in life old rose. Consistency hard, incompressible. | No ectosome. Choanosome: isodictyal reticulation; occasional irregular reticulation or tracts. | Oxeads: hastate, nearly straight, sharp points, 140–155 x 6–7 µm. |
| *Haliclona scabritia*<br>Tanita & Hoshino, 1989 | Japan | 62–72 | Thin, encrusting on stone, 50 x 35 mm x 1–2 mm thick. Surface punctate, smooth, almost even. Oscula not visible. Light pinkish beige in alcohol Consistency soft, fragile. | No ectosome. Endosome subisodictyal reticulation or occasionally irregular reticulation of oxeads. | Oxeads: uneven, straight to gently arched, rounded points, 130–170 x 8–9 µm. |
| *Haliclona tachibanaensis*<br>Hoshino, 1981 | Japan | Intertidal to subtidal | Irregular, thin to 1.5 cm with several swellings. Surface minutely hispid, uneven. Oscules 2–2.5 mm diameter, 0.8–1 cm apart on collars <1 mm high. Colour warm grey. Consistency slightly compressible, not tough. | No ectosome. Choanosome: reticulation of primary tracts and subisodictyal network. Primary tracts bi and trispicular, 15 µm diameter and ascending 50–120 µm apart. Primary tracts connected with subisodictyal reticulation. No tracts in deep part of sponge. | Oxeads: fusiform to hastate, straight to weakly bent in middle tapering to sharp points, 100–125 x 2–6 µm. |

| *Haliclona Species* | Loc | Depth (m) | External | Skeleton | Spicules |
| --- | --- | --- | --- | --- | --- |
| *Haliclona uwaensis*<br>(Hoshino, 1981) | Japan | Subtidal | Very thin encrusting. Oscula not visible. Colour ivory buff dry. Consistency very soft. | No ectosome. Choanosome: irregular coarse network of spiculo-fibre from 1–10 rows of spicules, 30 µm diameter. Numerous toxas and sigmas scattered in flesh | Oxeas: slightly arched, hastate, tapering to points, 150–185 x 610 µm. Toxas: thin, 28 µm. Sigmas: C, 10–22 µm. |
| *Haliclona takaharui*<br>Van Soest & Hooper, 2020 [*Haliclona viola* Hoshino, 1981] | Japan | Intertidal, low subtidal | Thin, 0.2–7 cm thick. surface smooth, uneven. Oscules 23 mm diameter, scattered over surface 0.7–1 cm apart. Oscula areas slightly swollen, 1–2 mm high. Colour in life deep purple. Consistency.soft. | No ectosome. Choanosome: vague tracts 10–30 µm diameter, 2–17 oxeas, ascending from substratum 50–100 µm apart. Cross connected with irregular reticulated oxeas. | Oxeas: hastate, straight or slightly bent at middle and sharp points, 130–155 x 5–8 µm. Oxeas: thin, fusiform, gently curved, tapering to points, 100–145 x 1–5 µm. |

Based on the comparisons discussed above *H*. (*R*.) *gesteta* is a new species and the only one recorded from BC which has not been reclassified from an original designation as *Reniera*.

### *Haliclona* (*Rhizoniera*) Griessinger, 1971

*Haliclona* (*Rhizoniera*) is defined as *Chalinidae with a regular anisotropic, ladder-like choanosomal skeleton consisting of pauci-to multispicular ascending primary lines, connected by unispicular secondary lines. Ectosomal skeleton usually absent, if present, consisting only of some vaguely strewn tangentially orientated oxeas. Spongin scarce or absent* (de Weerdt 2002 [2004]). Primary spicular tracts are typically somewhat wavy in BC specimens.

There are six species of *H.* (*Rhizoniera*) described for the North Pacific listed in Table 6; two of these are to subgenus only and two were misapplied to a Northeast Atlantic *Haliclona (Reniera) cinerea* (Grant, 1826).

**Table 6.** North Pacific *Haliclona* (*Rhizoniera*)

| Species | Location/Depth (m) | External | Skeleton | Spicules |
| --- | --- | --- | --- | --- |
| *Haliclona (Rhizoniera) enamela* Laubenfels, 1930 | Central California, intertidal | Encrusting to 2 mm thick with oscula on raised collars to 1.5 mm diameter. Colour in life is “drab”, i.e. light beige. | No ectosome; the choanosome has a reticulation of primary (6 to 8 spicules) and secondary (1 to 2 spicules) tracts. | Oscula 120–157 x 4.0–7.0 µm. |
| *Haliclona (Rhizoniera) rufescens* (Lambe, 1892 [1893])<br>Austin specimen: RBCM Cat. # 018-00370-001. | Lambe: Bering Sea, beach wash; Austin: Alaska, 0.13 m | Lobate subramose to nearly massive. Surface slightly rough to the touch. Oscula circular, each at the summit of a branchlet and with an average diameter of 2 mm. | Renierid in arrangement, a uniserial, moderately regular reticulation. | Oxeas: small, stout, rather sharply pointed, slightly curved, smooth; average size 144 x 13 µm. |
| *Haliclona (Rhizoniera)* sp. A (Hartman, 1975) (of Lee, et al. 2007) | Central California, intertidal and subtidal | Encrusting to 2 cm thick. Surface irregular. Oscules 2–7 mm in diameter, flush with surface or with raised rims to 10 mm high. Rose lavender | Vertical multispicular tracts, with cross spicules horizontal, diagonal, or irregular. | Oxeas: 130–188 x 3.0–11 µm. |
| *Haliclona (Rhizoniera)* sp. B (Hartman, 1975) (of Lee, et al. 2007) | Central California, intertidal | Encrusting to 5 mm thick. Surface smooth. Oscules less than 1–2 mm across, barely raised above surface. Grey-blue. | Vertical multispicular tracts; cross spicules horizontal, diagonal, or irregular. | Oxeas: 120–174 x 4.0–10 µm. |
| *Haliclona (Rhizoniera)* sp. (formerly-*Haliclona (Reniera) cinerea* (Grant, 1826) [of de Laubenfels, 1932]) | Central California, intertidal | Encrusting to 3 cm. Oscules conspicuous with raised, crater-like rims, 2-5 mm diameter. Surface, superficially very porous, crowded with depressions about 200 µm diameter. Lavender to drab brown. | Ectosome: inconspicuous. Fleishy dermal membrane. Choanosome: isodictyal reticulation with a few vague spicular tracts. | Oxeas: 135–169 x 4.0–9.7 µm. |
| *Haliclona cinerea* [of Lambe, 1893 [1894]] | BC, intertidal | Encrusting to 2 mm. 3 oscula 1 mm diameter. Dull yellow. | Not described. | Oxeas: 98–111 x 6 µm. |

*Haliclona (Rhizoniera) arborescens* n. sp. (Figure 8)

**Figure 8:**
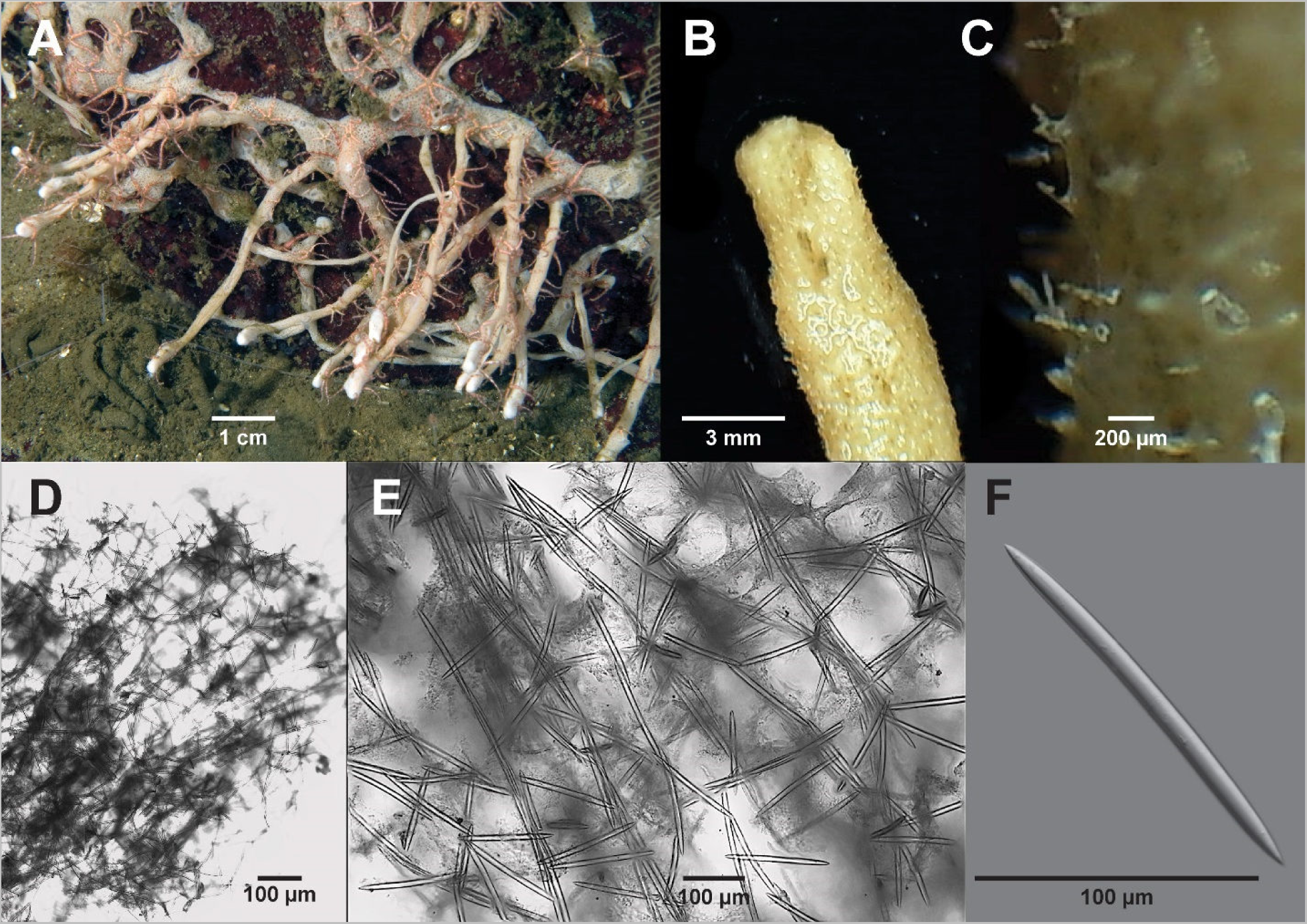
H*a*liclona (*Rhizoniera*) *arborescens* **n. sp.** A–F, Holotype, RBCM 018-00385-001. A in-situ. B Branch close up. C Branch surface. D Skeleton cross section. E Spicule tracts. F Oxea.

Zoobank yyyy

**Etymology** The name refers to the thin branching habitus of the sponge.

**Material Examined** Holotype RBCM 018-00385-001, station NM 415, 9 Mile Point, Sechelt Inlet, BC, 49° 36.304’ N / 123° 47.131’ W, 10 m depth, 8 May 2020, collector N. McDaniel, one specimen. Paratype RBCM 018-00133-001, station NM 317, Indian Arm, BC, 49° 25.134’ N / 122° 51.647’ W, 20 m depth, 14 Oct 2015, collector N. McDaniel, 1 specimen.

## Description

**External** (Figure 8A) Sponge flattened blind tubular branches that irregularly anastomose, spreading indefinitely; individual branches 6 x 0.3 cm on average. Surface microhispid (Figures 8B, C); spicules project individually up to 200 µm. Oscula not visible; ostia minute (preserved), densely covering sponge branches. Colour pale light brown to nearly white. Consistency: soft, easily compressed and fairly easily torn.

**Skeleton** (Figures 8D, E) No specialized ectosome. Skeleton composed of approximately parallel primary spicule tracts becoming vague in much of the sponge. Primary tracts are separated approximately 150 µm and are composed of 2 to 4 spicules varying in number along their lengths. Primary tracts are cross connected by single to double spicules predominantly approximately at right angles spaced at approximately 150 µm, forming a roughly isodictyal reticulation. Some meshes may be polygonal and the reticulation is not constant throughout the sponge body. Spongin absent.

**Spicules** (Figures 8F) Oxeas: gently curved, occasionally nearly straight; apices sharp, hastate, slightly mucronate, occasionally rounded. Rarely stylote. Immature oxeas fairly common, typically obtaining fully developed length before fully developed width rendering the apices acerate. Table 7 lists spicule dimensions of specimens examined (immature spicules excluded).

**Table 7.** *Haliclona* (*Rhizoniera*) *arborescens* **n. sp**. Oxea Dimensions (µm)

| Specimen | Length | Width |
| --- | --- | --- |
| NM 415<br>RBCM 018-00385-001 | 168 (204) 229 | 10.1 (11.4) 13.0 |
| NM 317<br>RBCM 018-00133-001 | 161 (198) 221 | 7.8 (10.4) 13.5 |

**Distribution** Two specimens have been found, widely separated: Sechelt Inlet (10 m depth) and Indian Arm of Burrard Inlet (20 m depth).

## Remarks

This sponge fits the subgenus *Haliclona* (*Rhizoniera*) due to the lack of spongin and lack of a specialized ectosome. Otherwise the skeletal architecture is more typical of subgenus *Haliclona*. See De Weerdt (2002 [2004]) for additional discussion. *Haliclona* (*R.*) *arborescens* **n. sp.** is branched unlike the other *H.* (*Rhizoniera*) species described in this report. There are currently 22 accepted species in the subgenus *Rhizoniera* (de Voogd, et al. 2024) with two reported for the North Pacific (Table 6). Referring to Table 6, none of the *H.* (*Rhizoniera*) species are branched except *H.* (*R.*) *rufescens* which has lobate branches but differs from *H.* (*R.*) *arborescens* **n. sp.** in having oscula on raised collars and smaller oxeas.

There are two *Haliclona* no subgenus reported for the North Pacific with a branching habitus: one from Japan, one from South Korea and one from the Pacific coast of Panama. *Haliclona daepoensis* (Sim & Lee, 1997) from South Korea has toxas as well as oxeas, *H. frondosa* Hoshino, 1981 from Japan has two sizes of oxeas. Based on these comparisons *H.* (*R.*) *arborescens* **n. sp.** is a previously undescribed species.

Haliclona (Rhizoniera) blanca n. sp. (Figure 9)

**Figure 9.**
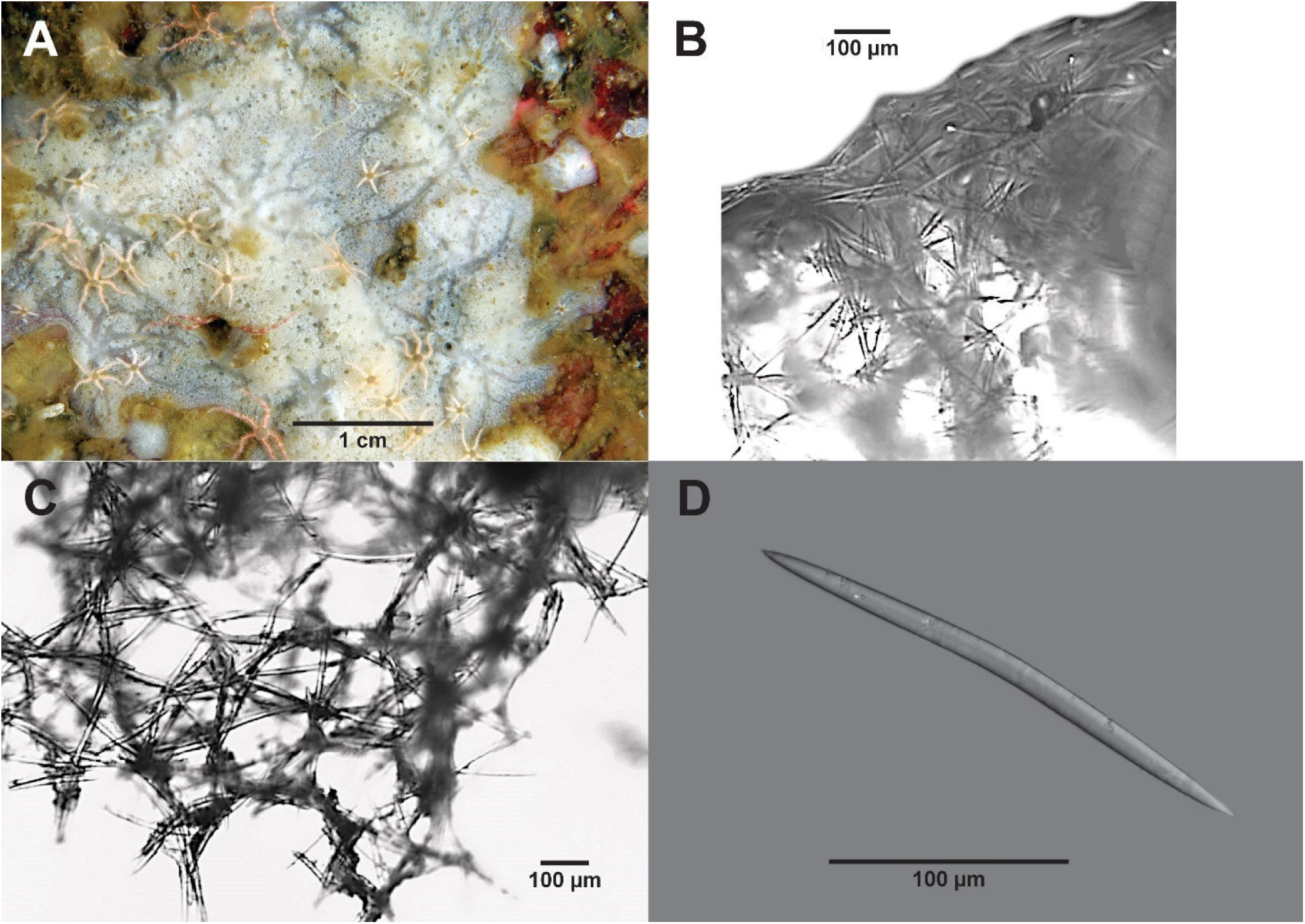
H*a*liclona (*Rhizoniera*) *blanca* **n. sp.** Holotype, RBCM 018-00152-008. A in-situ, B oblique view of ectosome, C choanosome, D Oxea.

Zoobank yyyyy.

**Etymology** The name refers to the colour of the living sponge.

**Material Examined** Holotype RBCM 018-00152-008, station NM 237, 9 Mile Point, Sechelt Inlet, BC, 49° 36.216’ N / 123° 47.396’ W, 23 m depth, 12 May 2011, collector N. McDaniel, G. Grognet, 1 specimen.

## Description

**External** (Figure 9A) Thin encrusting following contours of substrate, 3 cm diameter by 1.5 mm thick. Surface punctate with visible subdermal canals radiating from oscula. Oscula and pores both about 1 mm diameter. Colour in life white. Consistency soft, easily torn.

**Skeleton** Ectosome (Figure 9B) roughly tangential multilayer of oxeas 100 to 200 µm apart, 50 to 100 μm thick, penetrated in places by choanosome principal spicule tracts which project 50 to 100 μm. Choanosome (Figure 9C) fairly regular isodictyal reticulation 200 to 250 µm apart; principal multispicular tracts composed of two or more bundles cross connected by single spicules at approximately one spicule intervals.

**Spicules** (Figure 9D) Exclusively oxeas, most uniformly curved; some straight; most with moderately short apices: 208 (247) 268 x 9.1 (12.1) 14.3 µm. Styles rare and likely modified oxeas.

**Distribution** Known only from the type location: Sechelt Inlet, BC, 23 m depth.

## Remarks

The presence of a specialized ectosome sets this sponge apart from other northeast Pacific *Haliclona* species described in this report. None of the reported North Pacific *H.* (*Rhizoniera*) have a specialized ectosome (Table 6). *Haliclona cylindrica* (Tanita, 1961) accepted as *Haliclona tanitai* Van Soest & Hooper, 2020 from Japan is a cylindrical tube with a separate ectosome but spicules include toxas as well as oxeas. *Haliclona daepoensis* (Sim & Lee, 1997) from South Korea is an erect branching sponge with an ectosome of tangential oxeas and toxas. There are no other North Pacific *Haliclona* no subgenus with a specialized ectosome. Based on comparisons, *H.* (*R.*) *blanca* is a new species for the North Pacific.

*Haliclona (Rhizoniera) boothiensis* n. sp. (Figure 10)

**Figure 10.**
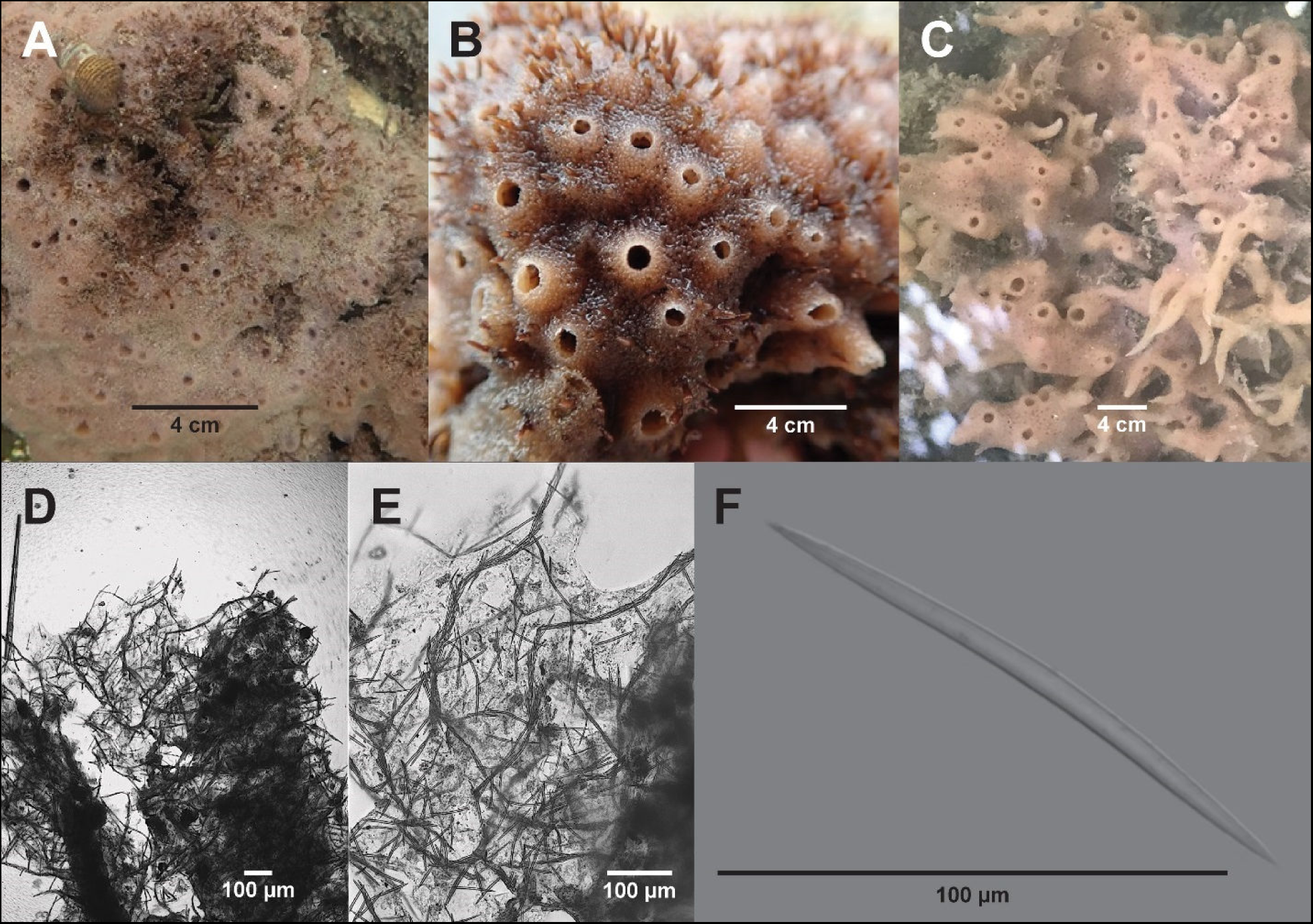
H*a*liclona (*Rhizoniera*) *boothiensis* **n. sp**. A, B, D–F Holotype, RBCM 019-00136-002. A in-situ. B close up in-situ. C variable habitus (photo P. Menning, DFO). D skeleton cross section at osculum. E Close up skeleton. F Oxea.

Zoobank yyyyy.

**Etymology** The name refers to the holotype location, Booth Inlet, Salt Spring Island, BC

**Material Examined** Holotype RBCM 019-00136-002, station RH 2019-09-13-04, Booth Inlet, Salt Spring Island, BC, 48° 51.868’ N / 123° 32.400’ W, intertidal, 13 Sep 2019, collector R. Harbo, 1 specimen.

Paratypes: RBCM 024-00003-001, station RH 2019-08-18-04, Booth Inlet, Salt Spring Island, BC, 48° 51.868’ N/123° 32.400’ W, intertidal, 18 Aug 2019, collector R. Harbo, 2 specimens. RBCM 024-00002-001, station RH 2019.09.13-01, Booth Canal Road, Salt Spring Island, BC, 48° 51.714’ N/ 123° 31.739’ W, intertidal, 13 Sep 2019, collector R. Harbo, 1 specimen. RBCM 024-00002-002, station RH 2019.09.13-02, Booth Canal Road, Salt Spring Island, BC, 48° 51.714’ N/ 123° 31.739’ W, intertidal, 13 Sep 2019, collector R. Harbo, 1 specimen. RBCM 018-00386-001, station RH 2000, Chain Point, Graham Is., Haida Gwai, BC, 53° 43’ N/132° 43’ W, intertidal, 28 Aug 2000, collector R. Harbo, 1 specimen. RBCM 024-00004-001, station RH 200725, Ayum Creek, Sooke Basin, BC, 48°23.433’ N/123°39.500’ W, intertidal, 25 Jul 2020, collector R. Harbo, 1 specimen. Station BO 18-08, Page Point, BC, 49^°^ 0.664’ N/ 123^°^ 49.241’ W, mid intertidal, 10 Sep 2018, collector R. Harbo, 1 non voucher specimen.

## Description

**External** (Figures 10A, B, C) Sponge encrusting, 7 x 12 cm x 5 mm thick (including chimneys). Surface densely packed with conical chimneys, 3 mm at base, 3 mm high with 1 mm diameter apical oscula. Some specimens have bent conical fistulae, blind or terminated by an osculum (Figure 10C). The area between chimneys is finely papillate. Colour in life pink to redish pink. Consistency spongy, easily torn.

**Skeleton** (Figures 10D, E) Branching and anastomosing multispicular tracts cross connected by single or multiple spicules at various angles forming a very irregular reticulation. Primary tracts are 1 spicule apart (about 100 µm) and 3–6 spicules wide. No specialized ectosome. Choanosome tracts carry to the surface and penetrate a few microns as single oxeas. Aquiferous canals numerous and elliptical (small forms) to elongate with curved sides (larger forms), 50 to 250 µm long axes.

**Spicules** (Figures 10E) Oxeas: curved, or straight, tapering from centre to acerate apices; a few immature; uncommonly centrotylote in some specimens. Oxeas 83–153 x 2.6–8.1 µm. Table 9 provides oxea dimensions for specimens examined.

**Table 9.** *Haliclona* (*Rhizoniera*) *boothiensis* **n. sp.** Oxea Dimensions (µm)

| Specimen | Length | Width |
| --- | --- | --- |
| **RH 2019-09-13-04**<br>RBCM 019-00136-002 | 104 (116) 143 | 4.4 (5.5) 7.5 |
| RH 2019-08-18-04<br>RBCM 024-00003-001 | 83 (100) 117 | 2.6 (4.7) 5.5 |
| RH 2019-09-13-01<br>RBCM 024-00002-001 | 101 (115) 130 | 5.2 (6.0) 7.8 |
| RH 2019-09-13-02<br>RBCM 024-00002-002 | 104 (119) 138 | 4.9 (5.8) 7.8 |
| RH 2000<br>RBCM 018-00386-001 | 78 (93) 104 | 5.2 (6.7) 8.1 |
| RH 200725<br>RBCM 024-00004-001 | 68 (96) 120 | 3.1 (6.3) 8.1 |
| BO 18-08 | 91 (104) 114 | 2.9 (5.4) 7.8 |

**Distribution** Haida Gwai, Salt Spring Island, Ladysmith Harbour, Hammond Bay and Sooke Basin (Aryn Creek), Vancouver Island, intertidal.

## Remarks

is similar to *Haliclona permollisimilis* Hoshino, 1981 in habitat (littoral), colour, oxeas and skeletal organization but differs in having oscula on chimneys. *Haliclona* (*Rhizoniera*) *boothensis* **n. sp.** is a fairly common intertidal *Haliclona* in BC waters. It is usually anchored to bedrock or large boulders and growing in association with green or red algae.

Haliclona (Rhizoniera) filix n.sp. (Figure 11)

**Figure 11.**
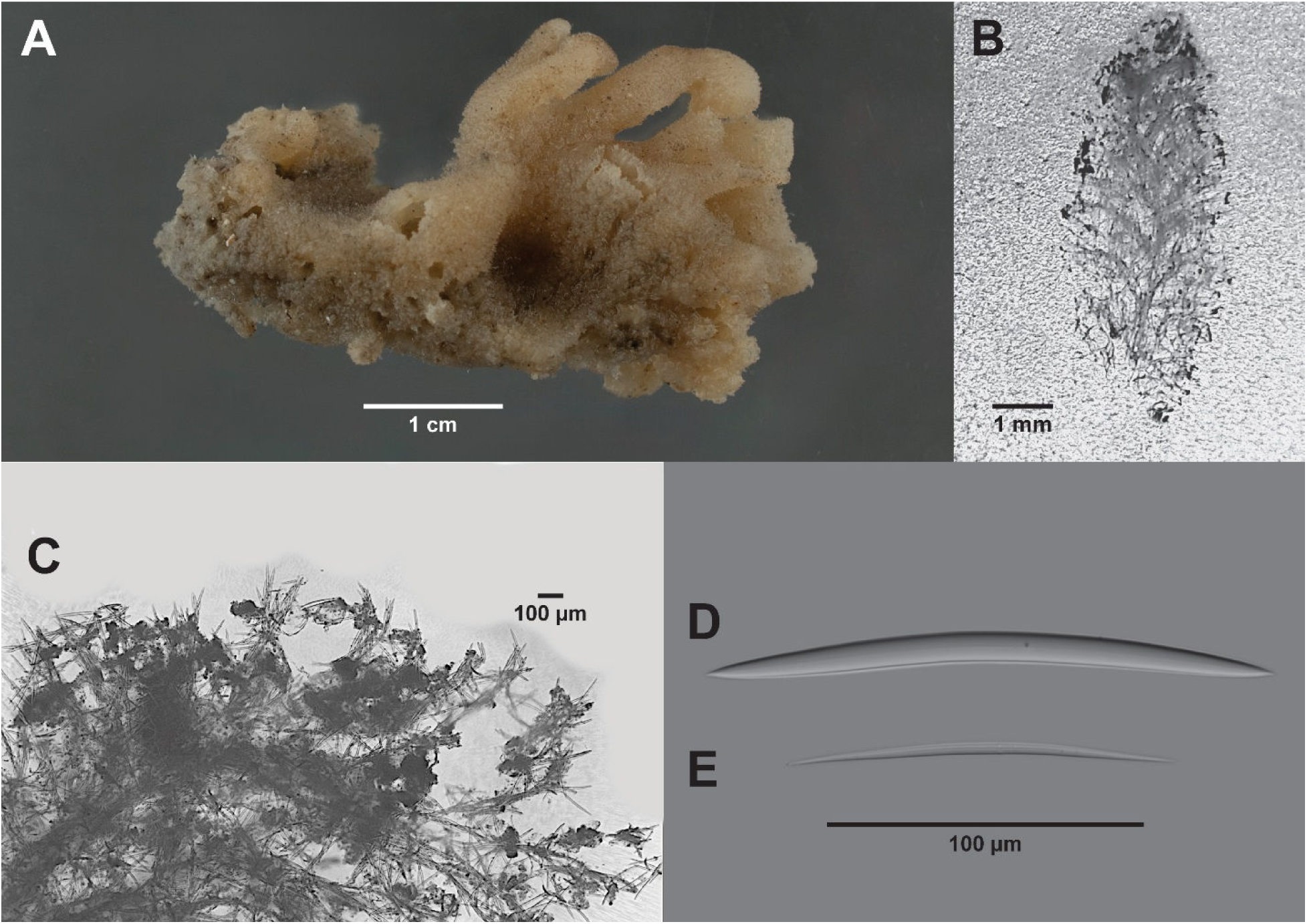
H*a*liclona (*Rhizoniera*) *filix* **n. sp.** Holotype RBCM 014-00161-03. A Preserved specimen. B Whole tube skeleton. C Close up near surface. D Fully developed oxea. E Immature oxea.

Zoobank yyyy

**Etymology** the species name is Latin for fern in reference to the fern-like appearance of longitudinal sections of the sponge tubes

**Material examined** Holotype RBCM 014-00161-03, station VT-81, Stuart Channel between Saltspring Island and Crofton, 48.865° N / 123.596° W, 34 m depth, 20 January 1981, collector V. Tunnicliffe, 1 specimen.

## Description

**External** (Figure 11A) Short tubes arising from a base. Tubes about 5–8 mm long x 2–3 mm diameter. Multiple tubes on a 2 x 3 cm base about 3 mm thick. Surface micropapillate.

Colour in alcohol beige. Consistency in alcohol, soft, compressible, easily torn.

**Skeleton** (Figures 11B, C) Skeleton of tubes forms a branching structure of principal tracts with a central tract and branches arching upward and outward to the surface where narrow brushes or plumes are formed about 2 spicules in length from the ends of the tracts. The stem tract and branches are 50–80 µm thick composed of 4 to 10 or more spicules across and spaced 100–200 µm apart except where aquiferous canals occur. At such locations, branches may be 300–400 µm apart. Single to sometimes multiple spicules cross connect the main tracts at irregular intervals from 50–200 µm forming a very irregular anisotropic reticulation. Spicule tracts in the base form an irregular reticulation similar to the tubes and carry up into the tubes. No spongin.

**Spicules** (Figure 11D, E) Spicules are exclusive oxeas, curved, slightly bent in the middle or straight; with acerate apices. The central canal of the oxeas is generally visible. Oxeas are 78 (93) 104 x 2.6 (5.7) 7.8 µm.

**Distribution** Known only from the Holotype location: Stuart Channel BC, 34 m depth.

## Remarks

The skeletal arrangement is peculiar for *Haliclona* (*Rhizoniera*) and sets this species apart from most of its congeners. Referring to Table 6, none of the North Pacific *Haliclona* (*Rhizoniera*) have oxeas as short as *H*. (*R*.) *filix* **n. sp.** and none form multiple thick tubes *H.* (*R.*) *rufescens* is lobate. However, *H*. (*R*.) *rufescens* has a renierid skeleton unlike that of *H*. (*R*.) *filix* **n. sp.**

The two *Haliclona* no subgenus species with sigmas are listed in Table 2; thin encrusting species are listed in Table 5. Branched species are discussed in the above section. *Haliclona daepoensis* (Sim & Lee, 1997), *H. shimoebuensis* (Hoshino, 1981), and *H. uwaensis* (Hoshino, 1981) have toxas; *H. liber* (Hoshino, 1981) has sigmas. *Haliclona cylindrica* (Tanita, 1961), *H. frondosa* Hoshino, 1981, *H. hoshinoi* Ise, 2017 [*H. punctata* Hoshino 1981 as *H*. (*Renaclona*) *punctata*], *H. sasajimensis* Hoshino, 1981, *H. ulreungia* Sim & Byeon, 1989, and *H. takaharui* Van Soest & Hooper, 2020 [*H. viola* Hoshino, 1981] have two sizes of oxeas. *Haliclona hydroida* Tanita & Hoshino, 1989, *H. robustaspicula* Hoshino, 1981, *H. sataensis* Hoshino, 1981, and *H. tachibanaensis* Hoshino, 1981 are thinly encrusting.

Of the remaining species *Haliclona bucina* Tanita & Hoshino, 1989 is infundibulaform and oxeas are larger than *H*. (*R*.) *filix* **n. sp**. (120–155 x 6–8 µm), *H. digitata* Tanita & Hoshino, 1989 is cylindrical and oxeas are larger than *H*. (*R*.) *filix* **n. sp**. (140–170 x 4–10 µm), *H. lentus* Hoshino, 1981 has a renierid skeleton without tracts, *H, permollisimilis* Hoshino, 1981, has only vague tracts in the skeleton and oxeas are larger than *H*. (*R*.) *filix* **n. sp**. (110–180 x 5–8 µm), and *H. sortitio* Hoshino, 1981 has oxeas larger than *H*. (*R*.) *filix* **n. sp**. (148–170 x 5–8 µm).

*Haliclona ellipsis* from Japan intertidal to shallow subtidal has oxeas slightly larger than *H*. (*R*.) *filix* **n. sp.** (90–120 x 4–6 µm) but has a renierid skeletal organization with only a few vague tracts. *Haliclona offerospicula* has small oxeas in the same size range as *H*. (*R*.) *filix* **n. sp.** (75– 90 x 2–3 µm) but is thin encrusting and has unispicular tracts. *Haliclona tenuis* has oxeas in the same size range as *H*. (*R*.) *filix* **n. sp.** (83–100 x 5–8 µm) but is very thin encrusting and has spicule tracts of two to three spicules wide, or thinner than *H*. (*R*.) *filix* **n. sp.**

Haliclona (Rhizoniera) kunechina n. sp. (Figure 12).

**Figure 12.**
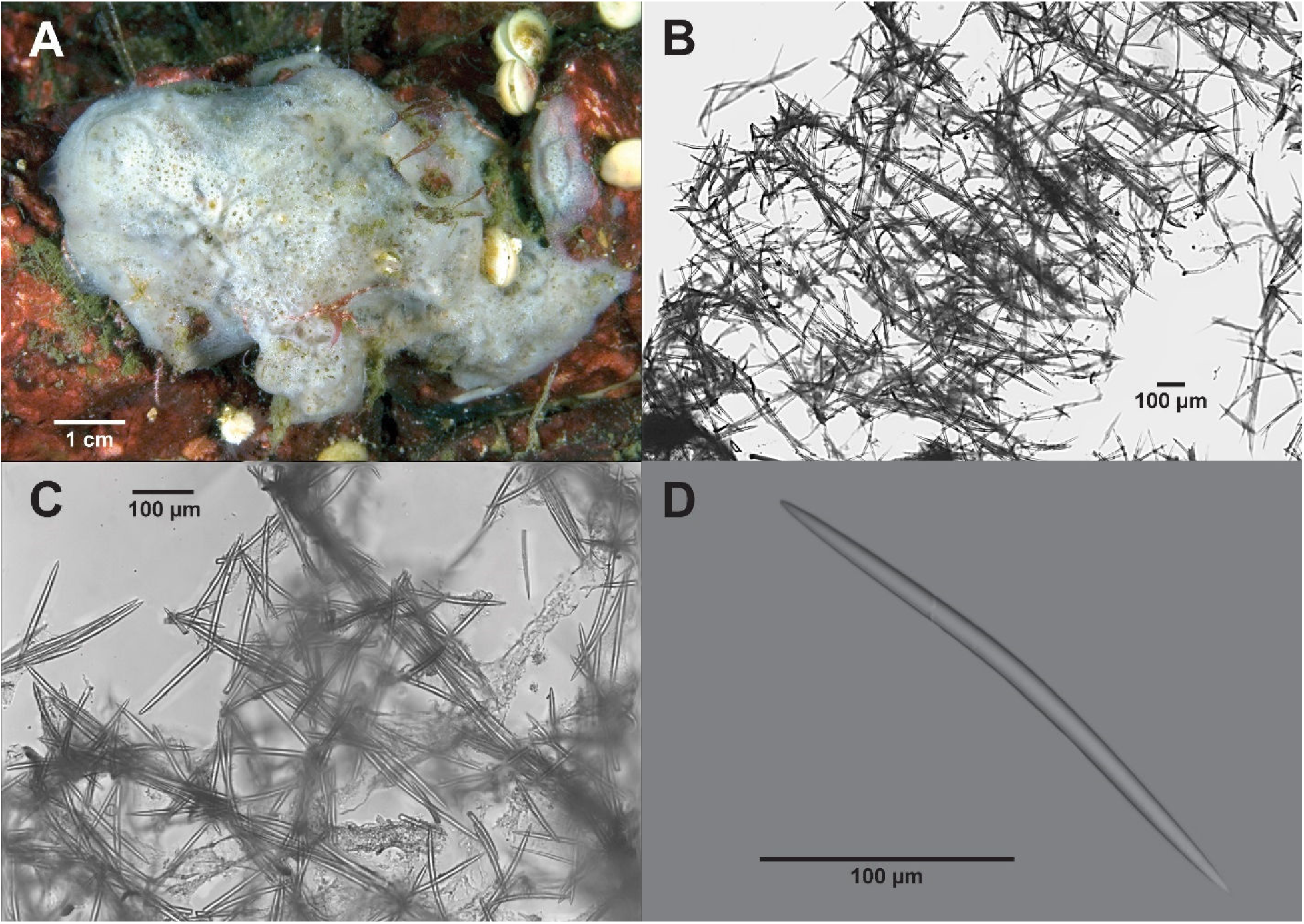
H*a*liclona (*Rhizoniera*) *kunechina* **n. sp.**, Holotype RBCM 018-00182-002. A in-situ. B Skeleton cross section. C Spicule tracts. D Oxea.

Zoobank yyyyy

**Etymology** Named for Kunechin Point, Sechelt, BC where the sponge was collected.

**Material Examined** Holotype RBCM 018-00182-002, station NM 279, Kunechin Point reef, Sechelt Inlet, 49° 37.060’ N / 123° 48.229’ W, 15 m depth, 8 Mar 2012, collector N. McDaniel, 1 specimen.

## Description

**External** (Figure 12A) Irregular, thickly encrusting 6 x 12 x 1 cm, very porous; numerous oscula slightly raised, approximately 2–3 mm diameter. Colour white. Soft, compressible, easily torn.

**Skeleton** (Figure 12B, C) No specialized ectosome. Multispicular tract cross connected by unispicular to a small number of spicules in an irregular reticulation. Multispicular primary tracts are 100 to 300 µm apart and may anastomose. Primary tracts are 20–30 µm thick.

Secondary spicules cross at right angles and are single or less commonly multispicular. Spongin only at some nodes. Many loose oxeas are disbursed among tracts.

**Spicules** (Figure 12D) Spicules are exclusive oxeas, slightly curved; occasionally straight. Apices variable, tending to hastate, slightly muconate, 216 (234) 260 x 7.8 (10.6) 13.3 µm.

**Distribution** Known only from the type locality, Kunchin Pt. reef, Sechelt Inlet, BC at 15 m depth.

## Remarks

*Haliclona* (*Rhizoniera*) *kunechina* **n. sp.** was full of diatoms and other micro phytoplankton and encrusting around green algal stems (green in the photo of the live sponge) but generally on bedrock. *Haliclona* (*Rhizoniera*) *kunechina* **n. sp.** oxeas at 216–260 µm are longer than reported North Pacific *Haliclona* (*Rhizoniera*) (Table 6). *Haliclona cylindrica*, *H. densaspicula*, *H. robustaspicula*, *H. shimoebuensis* and *H. violapurpura* have oxeas in about the same size range. *Haliclona cylindrica* has toxas as well as oxeas, *H. shimoebuensis* has two sizes of oxeas, *H. densaspicula* is very thin and has a hard texture, *H. robustaspicula* is thin encrusting and has an isodictyal reticulation, not spicule tracts, and *H. violapurpura* is a ramose sponge with a skeleton of subisodictyal reticulation and very vague tracts.

Haliclona (Rhizoniera) meandrina n. sp. (Figure 13).

**Figure 13.**
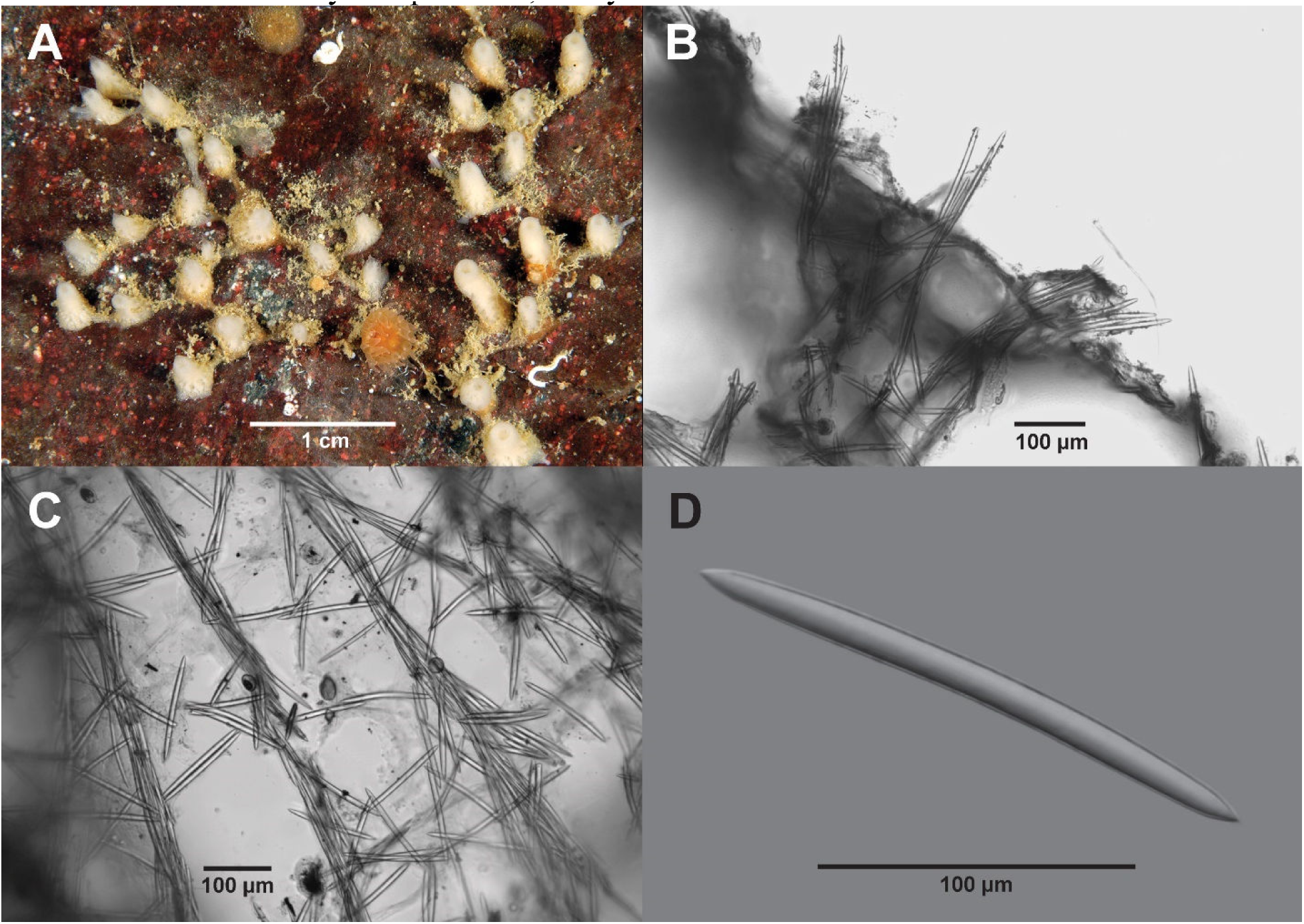
*Haliclona* (*Rhizoniera*) *meandrina* **n. sp.** A–D, Holotype RBCM 018-00171-003. A In-situ. B Ectosome. C Choanosome. D Oxea.

Zoobank *****.

**Etymology** The species name refers to the meandering habitus.

**Material Examined** Holotype RBCM 018-00171-003, station NM 259, Defence Islets, Howe Sound, 49° 34.544’ N / 123° 16.632’ W, 19 m depth, 19 May 2011, collector N. McDaniel, 1 specimen.

Paratypes: RBCM 018-00171-004, station NM 238, 9 Mile Point, Sechelt Inlet, 49° 36.216’ N / 123° 47.396’ W, 23.5 m depth, 2011-05-12, collectors N. McDaniel, G. Grognet, 1 specimen. RBCM 018-00171-005, station NM 251, Christie Islet, Howe Sound, 49° 29.919’ N / 123° 17.985’ W, 16 m depth, 2011.05.19, collectors N. McDaniel, D. Swanston, 1 specimen.

## Description

**External** (Figure 13 A) Branching and anastomosing; branches with conical tubes with oscula at the tips 1 mm diameter. Tubes 0.3 to 1 cm high by 0.3 cm diameter. Sponge spreading indefinitely, about 4 cm across. Surface microhispid (oxea project to 300 µm). Colour in life cream white. Consistency compressible, easily torn.

**Skeleton** Ectosome (Figure 13B) Strong primary ascending multispicular lines connected at irregular intervals by single or multiple spicules parallel tube long axis. Primary lines 1 to 2 spicule lengths apart; secondary lines 1 to 2 spicules long. Ascending and cross connecting tracts end at the surface in bouquets of a few oxeas more or less at right angles and penetrating the surface up to 300 µm; average about 100 µm. Exopinacoderm (dermal membrane) aspiculous or with scattered oxeas tangential to the surface. Large subdermal cavities beneath the exopinacoderm and between the bouquets. Choanosome (Figure 13C) anisotropic multispicular reticulation of oxeas. 1 to 5 spicules thick. Reticulation mostly quadrangular meshes. Length on side 1 spicule. Free spicules among the spicule tracts. Small amount of spongin at some nodes.

Central aquiferous canal leads to osculum at the apex of the cone. Choanosome fairly open with small to large aquiferous canals.

**Spicules** (Figure 13 D) Oxeas slightly curved, slightly mucronate hastate apices, 143–184 x 9.1–13.5 µm. Table 11 provides dimensions for the specimens examined.

**Distribution** Occurs in Sechelt Inlet and Howe Sound, BC.

## Remarks

None of the California *Haliclona* described by Lee et al. (2007) match *H*. (*R*.) *meandrina* **n. sp**. as detailed below:

- *Haliclona* sp. A & B of Hartman (1975) were not fully described by Hartman and placed in the subgenus *Rhizoniera* by Lee et al. (2007). None are conulose. *Haliclona* (*Rhizoniera*) sp., [formerly *Haliclona cineria* of de Laubenfels, 1932] has oscula with crater-like rims and a choanosome skeleton with few spicule tracts.
- *Haliclona* (*Haliclona*) *ambrosia* Dickinson, 1945 [of Sim & Bakus 1986] has cylindrical branches and a tangential skeleton.
- Other *Haliclona* identified only to genus, thus not fully characterized.
- *Haliclona* (*Haliclona*) sp. of Klontz 1989 as *Adocia* based on genus placement has a specialized ectosome. Oscula are on chimneys, not cones. Oxeas are shorter than those of NM 259 (120–140 µm vs. 143–235 µm).
- *Haliclona* cf. *permollis* (Bowerbank, 1866) [of Bakus & Green 1987] has sparsely spaced elevated oscula and oxeas 110–140 x 4–7 µm or at the lower range of NM 259. The sponge is purple, not white and oscula more scattered than NM 259.

*Haclona* (*Rhizoniera*) *rufescens* has lobate branches but a renierid not tracted skeletal architecture. *Haliclona* (*Rhizoniera*) *enamela* is encrusting and has a choanosome with spicule tracts. However, oscula are raised on collars and oxeas are generally shorter and thinner than *H*. (*R*.) *meandrina* **n. sp**. The habitus of *H*. (*R*.) *meandrina* **n. sp** is different from all of the *Haliclona* no subgenus species previously discussed.

Haliclona (Rhizoniera) penelakuta n. sp. (Figure 14)

**Figure 14.**
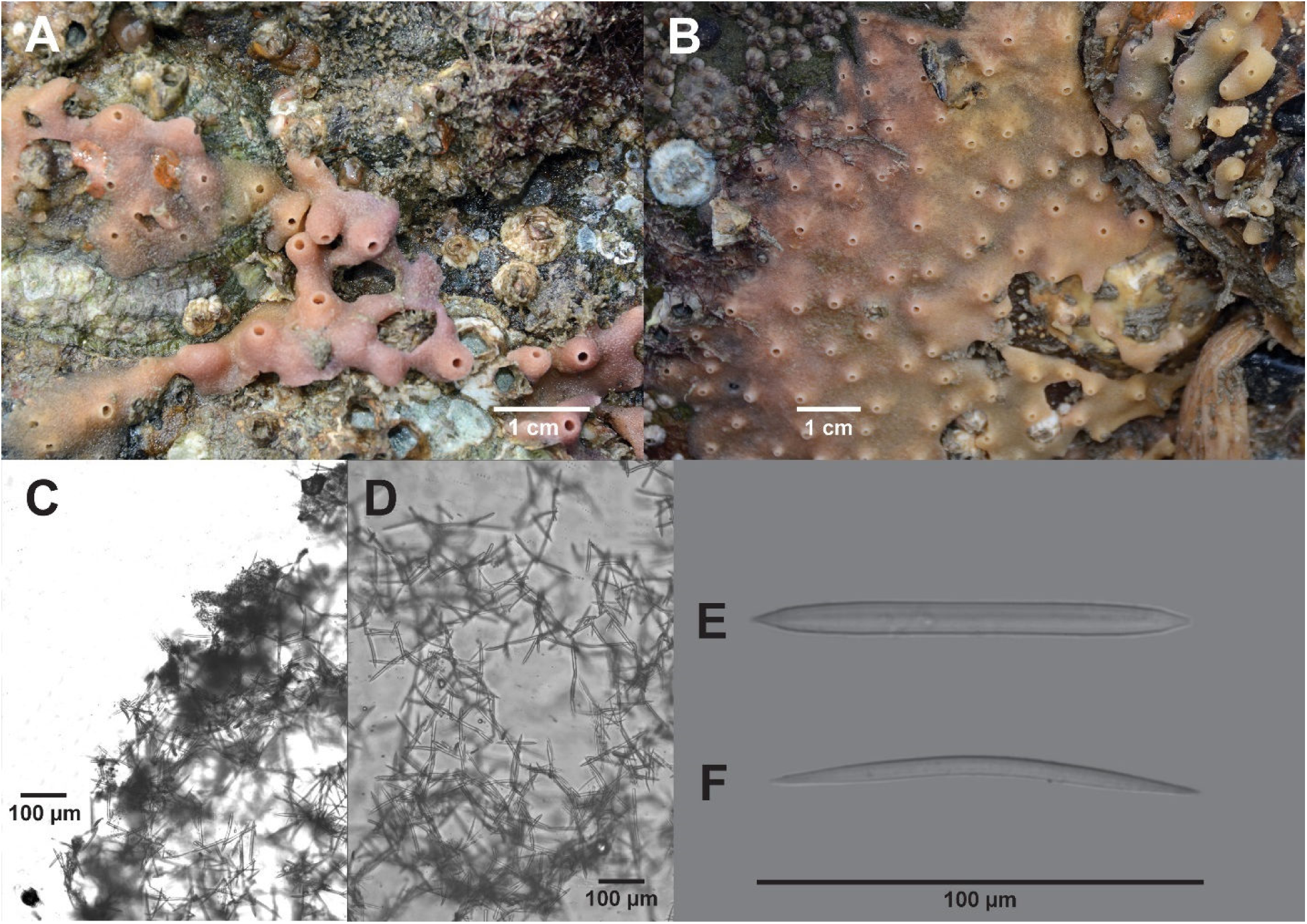
H*a*liclona (*Rhizoniera*) *penelakuta* **n. sp.** A, C–F, Holotype RBCM 018-00387-001. A meandering form. B spreading form. C skeleton near surface. D choanosome reticulation with large aquiferous canals. E thick oxea. F thin oxea.

Zoobank yyyyy

**Etymology** The name refers to the Holotype location, near Penelakut Island, BC.

**Material Examined** Holotype RBCM 018-00387-001, station NM 375, Penelakut Is., BC, 48° 58.957’ N / 123° 40.083’ W, intertidal + 2 m, 10 August 2018, collector N. McDaniel, 1 specimen.

Paratypes RBCM 018-00387-002, station NM 376, Penelakut Is., BC, 48° 58.957’ N / 123° 40.083’ W, intertidal + 2 m, 10 August 2018, collector N. McDaniel, 1 specimen. RBCM 018-00387-003, station NM 377, Penelakut Is. BC, 48° 58.957’ N / 123° 40.083’ W, + 2 m intertidal, 10 August 2018, collector N. McDaniel, 1 specimen. RBCM 018-00387-004, station NM 378, Penelakut Is. BC, 48° 58.957’ N / 123° 40.083’ W, + 2 m intertidal, 10 Aug 2018, collector N. McDaniel, 1 specimen. RBCM 018-00388-001, station NM 382, Boundary Bay, BC, 49° 02.30’ N / 123° 06.80’ W, +2-3 m, 23 August 2018, collector D. Swanston, 1 specimen; RBCM 024-00006-001, station Big Beach 2, Ucluelet BC, 48° 56.204’ N / 125° 33.124’ W, intertidal, 16 May 2022, collector R. Harbo, 1 specimen.

## Description

**External** (Figure 14A, B) Meandering to spreading laterally irregularly, encrusting oyster shells and bedrock to 2 mm. Surface micropapillate. Oscula raised on small cones 3 mm diameter at base and up to 3 mm high. Oscula 1–2 mm diameter. Encrustations 2–3 mm thick; 4– 5 mm thick at conulae. Colour flesh pink to yellowish brown. Consistency spongy, easily torn.

**Skeleton** (Figures 14C, D) No specialized ectosome. Subdermal spaces and somewhat more densely arranged spicules suggest an ectosome but spicule arrangement is as below the surface. At the surface spicules penetrate up to 100 µm at various angles singly, or in small clusters composed of up to 5 oxeas. Skeleton is an irregular anisotropic reticulation with vague wavy bi– and tri-spicular tracts, mostly in the lower parts of the sponge. Tracts 60–100 µm apart and 30–50 µm thick. The interior of the sponge is crossed by aqueous canals, the smaller roughly circular and 100 µm diameter. Larger canals are oblong and from 100 x 200 µm to 300 x 800 µm. The choanosome reticulation varies from triangular to rectangular to hexagonal. Loose spicules tend to obscure the reticulation. Spongin is sparse and only occurs at some reticulation nodes.

**Spicules** (Figures 14E, F) Oxeas: slightly curved or straight, long sharp (acerate) apices.

Thicker oxeas (ratio L:W>22.4) less common than thinner oxeas (ratio L:W<22.4). These “thick” oxeas are straight or slightly curved and have hastate mucronate apices. Stylote forms uncommon; immature forms fairly common in Holotype, but range in other specimens examined to uncommon. Table 10 provides spicule dimensions of specimens examined.

**Table 10.**
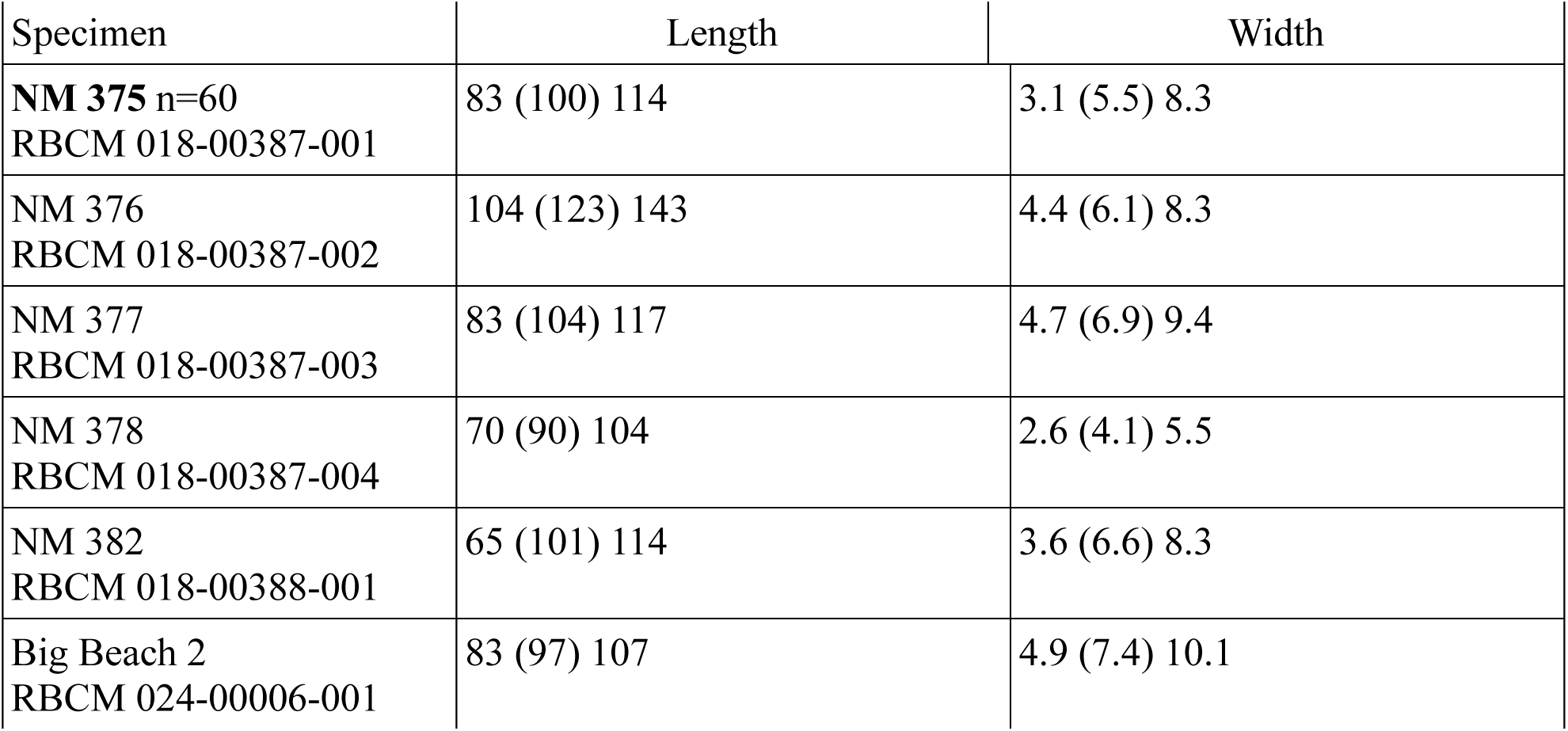
*Haliclona* (*Rhizoniera*) *penelakuta* **n. sp.** Oxea Dimensions (µm)

**Table 11.** *Haliclona* (*Rhizoniera*) *meandrina* **n. sp**. Oxea Dimensions (µm)

| Station | Length | Width |
| --- | --- | --- |
| NM 259<br>RBCM 018-00171-003 | 143 (180) 226 | 9.1 (11.8) 13.5 |
| NM 238<br>RBCM 018-00171-004 | 176 (213) 235 | 8.1 (11.4) 13.5 |
| NM 251<br>RBCM 018-00171-005 | 154 (169) 184 | 10.3 (12.0) 13.5 |

**Table 11.**
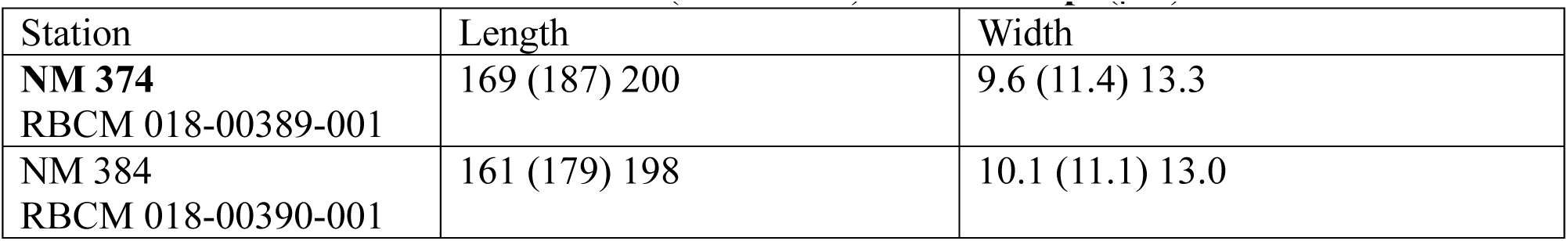
Oxea Dimensions of *Haliclona* (*Rhizoniera*) *vulcana* **n. sp**. (µm)

**Distribution** Between Thetis and Penelakut islands, in Boundary Bay, BC. and Ucluelet, BC. The sponge inhabits the high intertidal (+2–3 m depth) on both the inner and outer coasts of British Columbia.

## Remarks

Another Big Beach specimen (Big Beach 1) habitus and skeleton architecture are similar to the spreading form of *H*. (*R*.) *penelakuta* but the colour is mauve and the “thick” oxeas are absent. Referring to Table 6 none of the *H.* (*Rhizoniera*) species listed for the North Pacific has spicules as short as *H.* (*Rhizoniera*) *penelakuta* **n. sp.** except Lambe’s *H. cinera.* Lambe’s specimen had 1 mm diameter oscula and is possibly a close match. Lambe did not provide any details on the skeletal architecture of his specimen thus a further comparison is not possible. Referring to Table 5, *H. offerospicula* has spicules in the same size range but oscula are not on chimneys, the skeleton of *H. offerospicula* has unispicular tracts and the surface is smooth not microhispid as in *H.* (*Rhizoniera*) *penelakuta* **n. sp.**

Haliclona (Rhizoniera) vulcana n. sp. (Figure 15)

**Figure 15.**
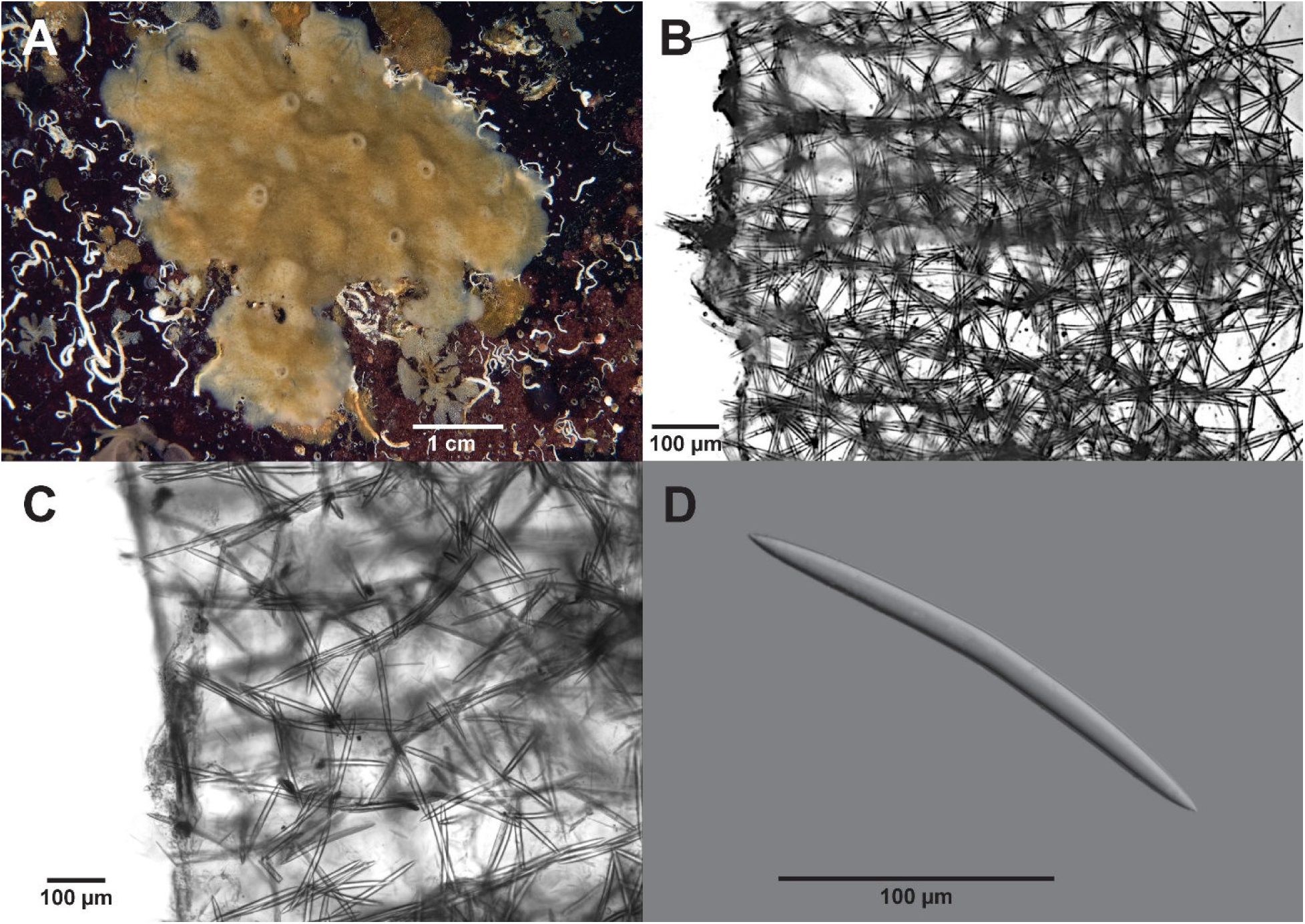
H*a*liclona (*Rhizoniera*) *vulcana* **n. sp.** A-D, Holotype RBCM 018-00389-001. A insitu. B Skeleton cross section. D Cross section near surface. D Oxea.

Zoobank yyyyy

**Etymology** Named for the volcano-like chimneys on the sponge.

**Material Examined** Holotype RBCM 018-00389-001, station NM 374, Escape Reef, Stuart Channel, BC, 48° 56.235’ N / 123° 39.430’ W, 15 m depth, 8 August 2018, collector N. McDaniel, 1 specimen. Paratype RBCM 018-00390-001, station NM 384, Grant Island, Welcome Pass, BC, 49° 30.662’ N / 123° 58.232’ W, 15 m depth, 2 March 2019, collector N. McDaniel, 1 specimen.

## Description

**External** (Figure 15A) Sponge 7 x 10 cm x 1.5 mm thick base. Chimneys 3 mm diameter at base x 3 mm high. Oscula 1.8 mm diameter. Ostia densely cover area between chimneys, 0.2 mm diameter. Surface rugose, microhispid; ridges radiate from some chimneys. Colour beige. Consistency easily torn.

**Skeleton** (Figure 15B, C) No specialized ectosome. Central part of choanosome with large (up to 0.8 mm diameter) aquiferous canals. Skeleton consists of approximately parallel bi– and tri-spicular primary tracts running vertically from the base to the surface. At the surface single spicules project up to 100 µm (Figure 15C). Primary tracts may branch and anastomose occasionally. Single spicules cross connect at right or random angles forming an irregular polygonal reticulation. Primary tracts 150–200 µm apart. Secondary spicules cross at 100 to 200 µm intervals. Spongin confined to nodes.

**Spicules** (Figure 15D) Oxeas: slightly curved or occasionally straight, sharp, hastate, slightly mucronate apices, A few thin immature oxeas present. Table 11 lists oxea dimensions of specimens examined.

**Distribution** Known from only two locations in southwestern BC: Stuart Channel and Grant Island, B.C. 15 m depth.

## Remarks

*Haliclona* (*Rhizoniera*) *enamela* habitus is similar to *H.* (*R.*) *vulcana* **n. sp.** but oxeas are shorter as are those of Lambe’s *H. cinerea*. The surrface of *H.* (*R.*) *enamela* is smooth and not microhispid. *Haliclona* (*Rhizoniera*) *rufescens* is lobate branched and the skeleton is renierid.

Hartman’s *H.* (*Rhizoniera*) species have flush oscula and oxeas are a bit shorter than *H.* (*R.*) *vulcana* **n. sp.** De Laubenfels’ (1932) *H. cinerea* oxeas are within the same size range as *H.* (*R.*) *vulcana* **n. sp.** Skeletal architecture is isodyctal reticulation and not tracted. Comparing North Pacific *Haliclona* no subgenus species listed in Table 5, *H. densaspicula*, *H. liber*, *H. scabritia*, and *H. uwaensis* have oxeas approximately in the same size range as *H.* (*R.*) *vulcana* **n. sp.**

*Haliclona* (*R.*) *densaspicula, H. liber* and *H. scabritia* have smooth surfaces not hispid and skeletons are renierid, not tracted; *H. uwaensis* has toxas in addition to oxeas.

## Discussion

*Haliclona* is a very successful genus. Worldwide there are over 470 species (de Voogd, et al. 2024) including subgenera and species not classified to subgenus. This is an increase of over 300 species in the genus *Haliclona* alone from the 150 extant species in the entire family Chalinidae (noted by de Weertd in 2000 [2004]) in a bit over 20 years enabled in part by creation of the World Porifera Database (www.marinespecies.org/porifera) and publication of Systema Porifera in 2002 [2004].

In some areas of the world where *Haliclona* occurs it is one of the most common sponges in the intertidal (de Weerdt 2002 [2004]) and this is the case in southwestern BC and California (de Laubenfels (1932), Lee et al. (2007)). Hoshino (1981) reports a similar abundance in Japan.

Species abundance and diversity in the Atlantic were detailed by Van Soest (1980) (southern Caribbean) and de Weerdt (1987, 1989, 2000 [2004]). This report places species names on a few of the more abundant *Haliclona* genera in southern BC waters and is a modest contribution to world *Haliclona* biodiversity.

Some *Haliclona* species are variable in colour and habitus as illustrated by the BC specimens examined. Colour variations occur in *Haliclona* (*Rhizoniera*) *penelakuta* **n. sp**. (Figure 14) and *Haliclona* (*Haliclona*) *mollis* (Lambe, 1893 [1894]) (Figure 5), a lavender –purple color to beige-brown. A number of BC *Haliclona* may also be brownish white to cream, e.g. *H.* (*Rhizoniera*) *arborescens* **n. sp.** (Figure 8), *H.* (*Rhizoniera*) *blanca* **n. sp.** (Figure 9), *H*. (*Rhizoniera*) *kunechina* **n. sp.** (Figure 12), *H*. (*Rhizoniera*) *meandrina* **n. sp**., (Figure 13), *H.* (*Rhizoniera*) *vulcana* **n. sp.** (Figure 15). Within-species habitus variations occur but are less frequent. It is well known the amount of current energy benthic invertebrates are exposed to effects habitus but some BC *Haliclona* species discussed vary within the same habitat. *Haliclona* (*Rhizoniera*) *penelakuta* **n. sp**. (Figure 14), from meandering ridges to spreading plates.

*Haliclona* (*Haliclona*) *mollis* (Lambe, 1893 [1894]) (Figure 5), from massive to various thickness encrusting. Because of their potential variability in colour and growth forms, they are difficult to identify in the field and microscopically as discussed below.

All subgenera of *Haliclona* were and are based on morphology (de Weerdt 2002 [2004], Van Soest 2017). The selection of subgenera of *Haliclona* is somewhat arbitrary since it depends (except for the subgenus *Gellius* with sigmas and the subgenus *Flagellia* with flagellosigmas) on skeletal architecture which often departs somewhat from the strict definition of the subgenus.

The subgenus *Haliclona* should always has spongin present (de Weerdt 1989, 2002 [2004]) but presence is variable and can also occur in other subgenera. Depending on the requirements for adherence to these definitions, specimens may be placed differently or not placed in a subgenus at all. The latter case appears to be particularly true of legacy species that are regarded as intermediate between subgenera or not described completely enough for subgenus placement. The genus has been shown by mitochondrial DNA and r-DNA analyses to be polyphylitic but at present morphology is generally relied upon to separate the subgenera (Bispo & Hajdu 2023), although some workers have reported success with sequencing of *Haliclona* (e.g. Muricy et al. 2015; Knapp, et al. 2015). We have chosen the subgenera for our new species which most closely fit the accepted skeletal architecture of the subgenera (de Weerdt 2002 [2004]) rather than adding to the list of *Haliclona* species not classified to subgenus. We have not attempted DNA analyses for this report.

